# Contributions of attention to learning in multidimensional reward environments

**DOI:** 10.1101/2023.04.24.538148

**Authors:** Michael Chong Wang, Alireza Soltani

## Abstract

Real-world choice options have many features or attributes, whereas the reward outcome from those options only depends on a few features/attributes. It has been shown that humans learn and combine feature-based with more complex conjunction-based learning to tackle challenges of learning in complex reward environments. However, it is unclear how different learning strategies interact to determine what features should be attended and control choice behavior, and how ensuing attention modulates future learning and/or choice. To address these questions, we examined human behavior during a three-dimensional learning task in which reward outcomes for different stimuli could be predicted based on a combination of an informative feature and conjunction. Using multiple approaches, we first confirmed that choice behavior and reward probabilities estimated by participants were best described by a model that learned the predictive values of both the informative feature and the informative conjunction. In this model, attention was controlled by the difference in these values in a cooperative manner such that attention depended on the integrated feature and conjunction values, and the resulting attention weights modulated learning by increasing the learning rate on attended features and conjunctions. However, there was little effect of attention on decision making. These results suggest that in multidimensional environments, humans direct their attention not only to selectively process reward-predictive attributes, but also to find parsimonious representations of the reward contingencies for more efficient learning.

**Significance Statement:** From trying exotic recipes to befriending new social groups, outcomes of real-life actions depend on many factors, but how do we learn the predictive values of those factors based on feedback we receive? It has been shown that humans simplify this problem by focusing on individual factors that are most predictive of the outcomes but can extend their learning strategy to include combinations of factors when necessary. Here, we examined interaction between attention and learning in a multidimensional reward environment that requires learning about individual features and their conjunctions. Using multiple approaches, we found that learning about features and conjunctions control attention in a cooperative manner and that the ensuing attention mainly modulates future learning and not decision making.

## Introduction

Every day, we face many choice options or actions that have a multitude of features or attributes. However, after making a decision, the feedback we receive is often binary (e.g., success or failure) or a simple scalar indicating how good or bad the outcome is, but not which feature(s) or attribute(s) resulted in the observed outcome. To guide decisions based on reward feedback, an agent could track reward history associated with the selection of each choice option, which can be seen as a unique configuration of multiple feature attributes. However, the number of possible combinations grows exponentially as the number of attributes grows, a problem referred to as the curse of dimensionality (Barto and Mahadevan, 2003; Sutton and Barto, 2018), making this strategy unfeasible due to memory constraints and insufficient reward feedback. Fortunately, in the real world, choice options that have similar perceptual properties also have similar reward values (e.g., most green fruits are unripe). Therefore, learning about features or attributes (i.e., feature-based learning) could provide an efficient strategy by avoiding learning about each choice option or action individually (Niv et al., 2015), thereby mitigate the curse of dimensionality without sacrificing much precision (Farashahi et al., 2017b).

In addition, feature-based learning allows the agent to deploy attention to process certain features more strongly during learning and/or decision making, resulting in additional flexibility (Mackintosh, 1975; Pearce and Hall, 1980; Dayan et al., 2000; Wilson and Niv, 2012; Niv et al., 2015; Akaishi et al., 2016; Soltani et al., 2016; Farashahi et al., 2017b; Leong et al., 2017; Oemisch et al., 2019). For example, attention can bias decision making by causing reward values associated with different feature dimensions to be weighted differently when they are combined. Moreover, attention can bias learning by attributing the reward outcome to certain feature dimensions, which leads the agent to preferentially update the values associated with those dimensions.

Unfortunately, feature-based learning becomes imprecise when certain features are predictive of reward values only when considered in conjunction with some other features (e.g., not all red or crispy fruits are edible but red crispy fruits usually are). Feature-based learning alone will ignore these important interactions, leading to incorrect generalizations. Of course, such imprecision can be mitigated by simultaneous learning about features and conjunctions of features (O’Reilly and Rudy, 2001; Farashahi and Soltani, 2021). However, this solution can become impractical as the number of conjunctions of features also grows exponentially. Importantly, not all feature conjunctions of a choice option are equally predictive of its reward value. Therefore, animals can achieve an appropriate performance if they also deploy attention to enhance learning about the most informative conjunctions of features and to prioritize those representations more strongly when making decisions. This necessitates a generalized form of selective attention beyond the selection among individual elementary features. It remains unclear whether this form of attention affects different types of learning strategies as well as choice behavior, and if so, how these effects interact with each other.

Finally, it is not fully understood how learned reward values influence attention on a trial-by-trial basis. In general, attention could be guided by reward value in multiple ways. For example, in some studies on simple and multidimensional reinforcement learning, attention has been shown to depend on the absolute difference (Hunt et al., 2014; Soltani et al., 2016; Farashahi et al., 2017b) or sum (Niv et al., 2015; Soltani et al., 2021), or maximum (Anderson et al., 2011; Leong et al., 2017; Gluth et al., 2018; Daniel et al., 2020) of the feature values of the alternative options. However, in most previous studies, the utilized reward schedules were simple, and these different functions on the values may lead to similar attentional weights, making it hard to elucidate the mechanisms by which reward value affects attention. Because all of these functions can be implemented through canonical neural circuits endowed with reward-dependent plasticity (Soltani and Wang, 2006, 2010), elucidating how value is transformed into attention can provide insight into the underlying neural mechanisms.

In general, an individual’s ability to use feature-based selective attention and apply configural learning (when required) falls under the umbrella of representation learning: to identify a compact but task-relevant representation of choice options that allows for efficient learning, including what feature is informative and how different feature dimensions interact (Niv, 2019; Radulescu et al., 2019a, 2021). Although previous studies have investigated different learning strategies separately, mechanisms of their interaction and how this interaction controls behavior are unknown. Understanding these processes provides an important step towards uncovering more robust solutions to the curse of dimensionality in complex, naturalistic environments.

Here, we used multiple methods including various reinforcement learning (RL) models with attentional components to examine human learning and choice behavior during a three-dimensional reward learning task to answer the following questions. First, how do simple feature-based learning and more complex conjunction-based learning interact to control choice behavior? Second, how do these learning strategies collectively determine or control attention (cooperatively or competitively)? Third, how and where do attentional modulations exert their influence: on choice, learning or, both? We use answers to these questions to discuss neural mechanisms by which attention shape representation learning in multidimensional reward environments.

## Materials and Methods

### Participants

In total, 92 healthy participants (N=66 females) were recruited from the Dartmouth College student population (ages 18–22 years). Participants were recruited through the Department of Psychological and Brain Sciences experiment scheduling system at Dartmouth College. They were compensated with money and T-points, which are extra-credit points for classes within the Department of Psychological and Brain Sciences at Dartmouth College. All participants were compensated at $10/hour or 1 T-point/hour. They could receive an additional amount of monetary reward for their performance of up to $10/hour. Similar to a previous study based on the same dataset (Farashahi and Soltani, 2021), we excluded participants whose performances (proportion of trials in which the more rewarding option is chosen) after the initial 32 trials were lower than 0.53. We also excluded an additional participant who failed to provide any reward probability estimates in 3 out of 5 bouts. These criteria resulted in the exclusion of 25 participants in total. All experimental procedures were approved by the Dartmouth College Institutional Review Board, and informed consent was obtained from all participants before the experiment.

### Experimental paradigm

The multidimensional reward learning task involved learning about the reward values (reward probabilities) associated with multidimensional visual stimuli through reward feedback and consisted of choice and estimation trials. Stimuli consisted of three feature dimensions (color, shape, and texture) where each feature dimension had three possible values (three colors, three shapes, and three textures), leading to 27 stimuli (objects) in total. During the choice trials, the participants were presented with two stimuli that had distinct features in all three dimensions, and they were asked to choose between them to obtain a reward. Reward feedback (whether they won a reward point or not) was provided randomly after each choice with a probability determined by the reward schedule. In choice trials, the order of the stimuli’s presentation was pseudo-randomized such that all pairs that were distinct in all three feature dimensions were presented four times in different spatial layouts. During the estimation trials, participants were presented with each of the 27 stimuli in random order and were asked to estimate the probability that the selection of each stimulus would lead to a reward. There were 432 choice trials in total. The estimation trials were interspersed in five bouts that appeared after choice trial numbers 86, 173, 259, 346, and 432.

The reward schedule (*p*_r_(*O*), ∀ stimuli/objects *O*) determined the probability that selection of a given stimulus/object was followed by a reward. We used a reward schedule with a moderate level of generalizability (Farashahi et al., 2017b) such that one feature dimension (informative feature) and the conjunction of the two other features (the informative conjunction) were predictive of reward on average to some extent. For simplicity, we used *F_i,j_* to denote the *j*th instance of the *i*th feature, where *i* = {1, …, *m*}, *j* = {1, …, *n*}. We also used *F_k_*(*O*), *C_l_*(*O*) to denote the *k*th feature and the *l*th conjunction of stimulus/object *O*, respectively. Finally, the *l*th conjunction denoted the conjunction of the two features other than feature *l*.

An algorithm analogous to the Naive Bayes algorithm can be used to approximate the reward values associated with each stimulus using the average reward probabilities associated with its features, as detailed below (Murphy, 2012; Hunt et al., 2014; Farashahi et al., 2017b), under the simplifying assumption that individual features are conditionally independent given the reward outcome. The average reward probability for different instances of a given feature can be obtained by marginalizing over other feature dimensions. We use 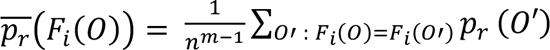 to denote the value of the *i*th feature of the stimulus/object *O*. Given these *feature values*, the reward probability of stimulus/object *O* can be estimated as follows:

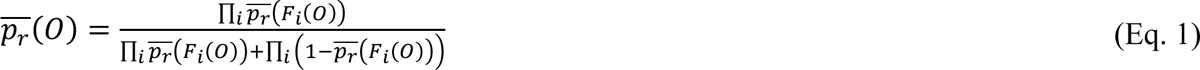

Therefore, the log-odd of reward for stimulus/object *O* is a linear combination of the log-odds of average reward probabilities for its features, estimated using the marginalization method calculated above: 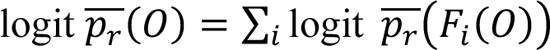 (Hunt et al., 2014; Farashahi et al., 2017b). A similar process could be defined for estimating stimulus values based on a mixture of *feature* and *conjunction values*:

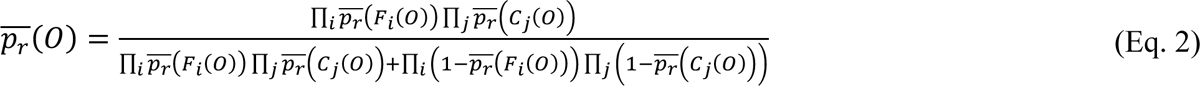

where we use 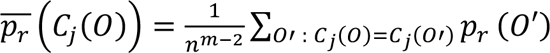 to denote the value of the *j*th conjunction of the stimulus/object *O*. Because the conjunctions *C_inf_* should not contain any of the features *F_inf_*, there are different ways of combining features into conjunctions. In the case of three features as in the current study, there are three ways of using mixed feature and conjunction values to represent the stimulus/object value. Different strategies can lead to different amounts of approximation error, which we define as the average KL divergence between *p*_r_(*O*) and *p*_r_(*O*) for all objects *O*. The more features are merged into conjunctions, the more accurate the estimation strategy can be with enough data, but the costlier it is to calculate. For the same reason as above, 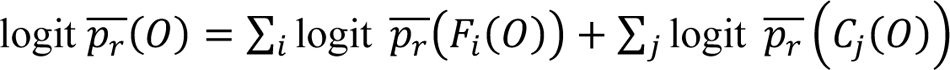 (Farashahi and Soltani, 2021). The above formulae present the optimal combination of feature values, or feature and conjunction values. In reality, however, estimated reward values might be a weighted sum of the feature and conjunction values.

### Computational models of value-driven attention

To capture participants’ trial-by-trial learning and choice behavior, we fit reinforcement learning models that estimated different types of reward values (i.e., probability of reward associated with features, conjunctions, or stimuli) and included different types of attentional modulation during choice and decision making. To test how learned reward values drive attention, we compared three possible relationships between the reward values of the two presented options and attentional modulation: uniform attention (no relationship), attention based on summed values, attention based on the absolute difference in values, and attention based on the maximum value.

In addition, we assumed that attention could modulate choice, learning, or both, which led to a total of ten variations of attentional mechanisms. We also considered five different *learning strategies* (1) *F*: feature-based learning, (2) *F* + *O*: feature- and object-based learning, (3) *F* + *C_Separate_*: feature- and conjunction-based learning with independent attention, where attention weights on the features and conjunction are calculated separately, (4) *F* + *C_feature attn_*: feature- and conjunction-based learning where attention only biases features and is uniform over conjunctions, and (5) *F* + *C_joint_*: feature- and conjunction-based learning, where attention is based on the integrated values of features and conjunctions, and modulates both features and conjunctions. We tested the last three variations of learning strategies to investigate possible mechanisms of interaction between attention based on feature values and conjunction values.

This led to 50 models in total (5 learning strategies × 10 attentional mechanisms).

We based all our learning models on the RL models with decay, which have been shown to capture behavior in similar tasks successfully (Niv et al., 2015; Farashahi et al., 2017b; Farashahi and Soltani, 2021). This means that while the values associated with features, conjunctions, and/or the object identity of the chosen option were updated after each reward feedback (*r*(*t*) = 0 or 1), all other values decayed towards a baseline (see below). All the values were initialized at 0.5, and values associated with the chosen options (*V_ch_*) were updated after each feedback using the following equations:

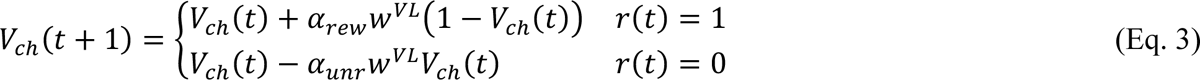

where *w^VL^* is the attentional weight used to modulate value learning, α*_rew_* and α*_unr_* are the learning rates for rewarded and unrewarded trials, and *ch* stands for the feature, conjunction, and object index associated with the chosen option. Based on results from previous studies, we set the baseline for the decay of the unchosen (and unavailable options) values (*V_unch_*) to 0.5 as follows:

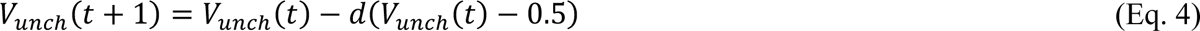

This decay process was not modulated by attention.

The log odd for choosing a stimulus on the left was the weighted sum of feature, conjunction, and/or object values of that stimulus as follows:

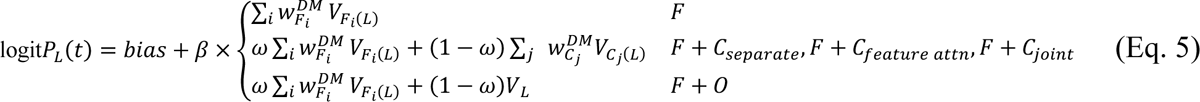

where logit*P* = log*P*/(1 − *P*), *w*^DM^ are the attentional weights used to modulate decision making, ω is a parameter that interpolates between feature- and conjunction (or object)-based learning, β is the inverse temperature parameter, and *bias* is the bias towards choosing the option on the left. The attentional weights were calculated based on the following equations:

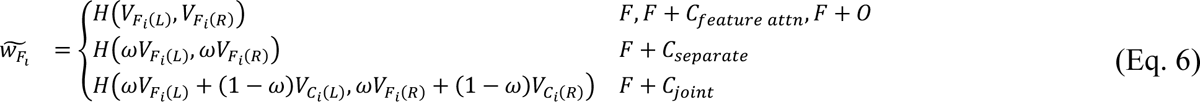

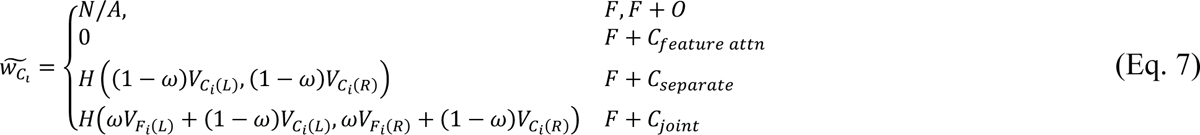

where the function that combines the reward values of alternative options on each trial could be *H*(*x*, *y*) = 0 for uniform weighting, *H*(*x*, *y*) = (*x* + *y*)/2 for sum or average weighting, *H*(*x*, *y*) = |*x* − *y*| for absolute difference, or *H*(*x*, *y*) = max(*x*, *y*) for maximum value. The attention weights for features (and analogously for conjunctions) were normalized through a softmax function:

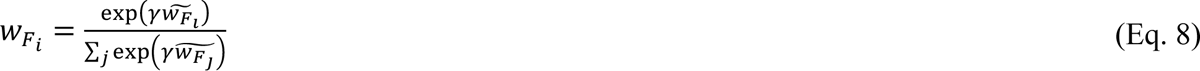

This resulted in a maximum of 7 parameters for the tested models: *bias*, β, ω, *d*, α*_rew_*, α*_unr_*, and γ.

### Model fitting and model selection

The RL models were fit using the Bayesian Adaptive Direct Search (BADS) optimization algorithm (Acerbi and Ma, 2017). Forty random initial optimization points were sampled to avoid local optima. For each model with attention, one set of the initial parameter values was chosen as the best parameters for the base model with the same learning strategy but without attention. Models were compared using random effects Bayesian Model Selection (BMS) based on the Bayesian Information Criterion (BIC) as an approximation for model evidence (Stephan et al., 2009; Rigoux et al., 2014). We report the posterior model probability, which is the posterior estimate of a model’s frequency as well as the protected exceedance probability (pxp), which equals the probability that one model exists more frequently than all other models. Unless otherwise specified, the Bayesian omnibus risk (BOR), which measures the probability that the observed differences in model frequencies are due to chance, was less than 0.001 for all our model comparisons. This suggests that there were significant discrepancies in different models’ ability to account for behavioral data.

For all mixed-effects modeling, we fit a random slope and intercept for each participant. If the model did not converge, we incrementally simplified the random effect structure by first removing correlation between random effects, followed by the random slopes. If the model failed to converge even with only random intercepts, an ordinary linear regression model was used.

### Model-based analysis of choice behavior

Using the estimated model parameters from fitting the choice data with the best model, we were able to simulate our models and examine the latent variables (Wilson and Collins, 2019), including the inferred subjective values and attentional weights. We computed the entropy of the attentional weights 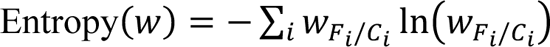 to examine how concentrated the attentional weights were on each trial. Lower entropy means more concentrated attention and a stronger bias towards a single feature or conjunction. We also used the Jensen-Shannon divergence (JSD) between the attentional weights at consecutive trials,

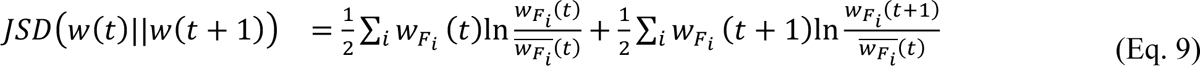

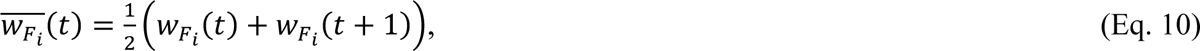

to characterize changes in attention between trials. The higher the JS divergence (JSD), the larger the change in attention between trials (JSD is bounded between 0 and 1). We also used differences in trial-wise BIC of different models to measure transition between different learning strategies (Farashahi et al., 2017b; Farashahi and Soltani, 2021), where trial-wise BIC is defined as

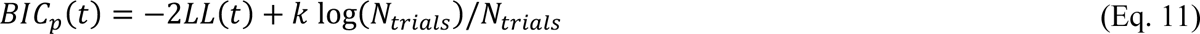

We used linear mixed-effects models to predict these quantities as a function of trial number (with trial number normalized between 0 and 1) to characterize their time courses.

### Model-free analysis of choice behavior and value estimation

To identify biases in reward credit assignment and choice, we fit generalized linear mixed-effect models to predict choice based on reward and choice history early in the experiment (first 150 trials) and based on objective reward values at the end of the experiment (when reward assignments have been materialized) as described below.

To examine bias in reward credit assignment, we generalized win-stay lose-switch commonly used to study learning in simple environments (Lau and Glimcher, 2005; Noonan et al., 2010, 2017; Walton et al., 2010; Katahira, 2018; Moran et al., 2019) to our multidimensional environment. More specifically, we used the following generalized linear mixed model (GLMM):

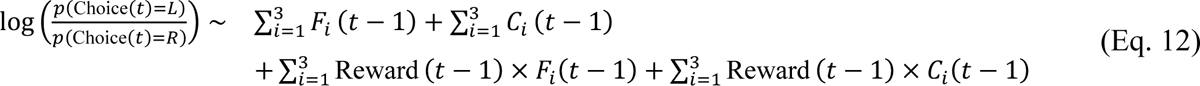

where *F*_i_(*t* − 1) and *C*_i_(*t* − 1) were equal to +1 if the option on the left shared the *i*th feature or *i*th conjunction as the previous chosen option, −1 if the option on the right shared the *i*th feature or *i*th conjunction as the previous chosen option, and 0 if neither options shared the feature or conjunction with the previously chosen option. Reward (*t* − 1) was equal to +1 and −1 if the previous choice was rewarded and unrewarded, respectively. The first two terms capture the tendency to repeat choosing the same feature or conjunction, regardless of reward feedback, whereas the last two terms capture the tendency to repeat or avoid a previously chosen feature or conjunction depending on reward feedback (generalization of win-stay and lose-switch).

We used the first 150 trials to study reward credit assignment because participants’ performances reached their steady state at that point, and because participants’ sensitivity to reward history should decrease with more learning. Although the GLMMs cannot separate the influence of attentional modulations at the time of choice and learning (Katahira, 2018), they can nonetheless detect biases in choice behavior. Importantly, attention would lead to some coefficients being higher than the other ones, indicating heightened sensitivity to the reward history associated with some feature(s) and/or conjunction(s). We note that choice auto-correlation could be due to a tendency to choose options that share a feature or conjunction with a past choice, or due to biased learning that weighs positive outcomes more heavily than negative outcomes. Although both mechanisms may be at play (Katahira, 2018; Palminteri and Lebreton, 2022), this second explanation implies that when there is positivity bias in learning, choice auto-correlation could also be biased by attention, leading to increased sensitivity to the choice history associated with some feature(s) and/or conjunction(s).

To examine the participants’ choice strategy at the end of the experiment, we fit GLMMs that used the values associated with the informative feature and the informative conjunction to predict participants’ choice. If the participants employed a feature- and conjunction-based learning strategy, the values associated with both the informative feature and informative conjunction should significantly predict choice.

Finally, to examine participants’ value representations more directly, we also fit their estimations of the reward probability associated with each stimulus/object with GLLMs that used reward values along different feature or conjunction dimensions as the independent variables. Both the reward probability estimates and the marginal probabilities were transformed through a logit transformation log *p/1-p*. We fit five models using the following independent variables, (1) *F_inf_*: the value of the informative feature, (2) *F_inf_* + *C_inf_*: weighted sum of the values of the informative feature dimension and the informative conjunction, (3 and 4) *C_noninf1_*, *C_noninf2_*: the values of the two non-informative conjunctions (the marginal probabilities along the two non-informative features are either all 0.5 or between 0.48 and 0.52, therefore too close to 0.5 to be meaningful), and (5) *O*: actual reward probabilities associated with each object/stimulus. Due to the high correlation between reward values of the informative feature and reward values of the non-informative conjunctions (*R* > 0.9), we were not able to include those in the same model.

Instead, we fit separate models using reward values along different stimulus dimensions and compared the different models’ goodness-of-fit. To avoid biases from using predefined marginal probabilities and account for any distortions of learned values, we also fit mixed-effects ANOVA models using each of the three features as a factor and included all interaction terms. This allowed us to investigate how much variance in the value estimations is explained by each dimension by looking at the partial eta-squared (η_p_^2^) statistic. Because during each trial, each participant only gives one estimation for each object, we only include random effects up to two-way interaction terms.

### Model recovery and model validation

Due to the large number of models we used to fit the data, a comprehensive model recovery was intractable. Instead, we performed a model recovery of the best-fitting model against other models that share the same learning strategy or the same attentional mechanism. To perform model recovery, we used the same reward schedule, and sampled 100 stimuli sequences using the same rule as for participants in the experiment. For each model type, we sampled parameter sets uniformly or log-uniformly from a plausible range of values. Consistent with the experiment, we kept only parameters that produced a choice sequence with a performance better than our exclusion threshold. We also removed simulated trials that produced highly random choice sequences (choice sequences that had an average likelihood per trial of less than 0.525 given the sampled parameters). Based on these parameters, we simulated choice sequences using the actual sequences of stimuli observed by each participant. We then fit all models and compare them using random effects Bayesian model selection. We also verified parameter recovery by reporting Spearman’s rank correlation between the ground truth parameters and the best fit parameters.

Finally, to qualitatively validate our winning model, we used the estimated parameters based on the best model to simulate choice sequences and trial-by-trial attention weights. We then examined qualitative match between the experimental and simulated data.

## Results

To investigate the effects of attention on learning and decision making in high-dimensional environments, we re-analyzed choice behavior in a multidimensional probabilistic learning (MDPL) task (Farashahi and Soltani, 2021) in which human participants selected between pairs of visual stimuli (objects), each defined by three visual features (color, pattern, and shape). Each feature could take three values, making a total of 27 stimuli or objects to learn about. Each choice was followed by a binary reward feedback with the probability of harvesting a reward determined by the reward schedule. Participants learned about these reward probabilities through trial and error (see Materials and Methods for more details). We also asked participants to provide their estimations of reward probabilities for each stimulus at five evenly spaced time points throughout the experiment (**Fig. 1A**). Critically, the reward probability associated with the selection of each stimulus was determined by the combination of its features such that one informative feature and the conjunctions of the other two non-informative features (the informative conjunction) could accurately predict reward probabilities (**Fig. 1C**). Nonetheless, reward probabilities associated with the 27 stimuli greatly varied according to the presence of the informative or non-informative features (**Fig. 1D**), making it a non-trivial task to find the informative feature, unlike previous experiments (Niv et al., 2015; Leong et al., 2017; Oemisch et al., 2019). By learning about and combining the predictive values of the informative feature and conjunction, participants could achieve a good approximation to the actual reward probabilities associated with each stimulus. In contrast, learning about the non-informative feature and conjunction dimensions leads to inaccurate approximation while still allowing performance that was better than chance (**Fig. 1E**; see also Experimental Paradigm in the **Materials and Methods**).

**Figure 1.**
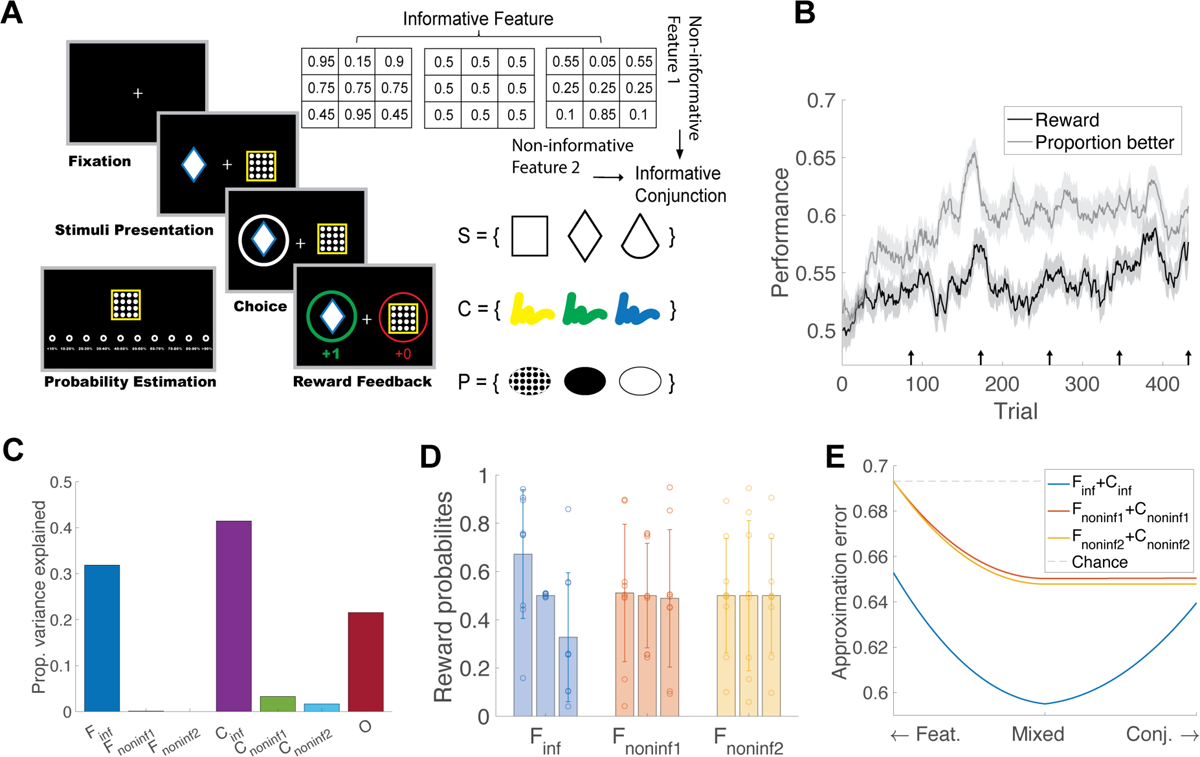
The experimental design and overall performance. (**A**) The timeline of each choice trial consisting of fixation, stimulus presentation, choice presentation, and reward feedback. Five bouts of probability estimation trials were interleaved throughout the 432 choice trials. The reward probabilities of the 27 stimuli, each identified by 3 visual features, are shown in the inset. (**B**) The average learning curve across all participants. The black and gray curves show the average reward received and the proportion of trials the participants chose the better option, respectively. Both curves were smoothed by a moving average filter over 20 trials. The shaded area indicates the SEM. Performance reached the steady state at around 150 trials. Arrows on the x-axis indicate the trials after which a value estimation trial was administered. (**C**) Informativeness of each dimension as measured by the proportion of variance in the reward schedule explained by each feature, conjunction, and the stimulus (object) identity. The informative feature and conjunction predict a larger amount of variance in the reward schedule than non-informative ones. Additional variance can only be explained by the stimulus (object) identity. (**D**) The average reward value of individual stimuli ordered by each feature dimension and contained feature. The height of the bars shows the mean values with error bars indicating standard deviation, and circles show the exact stimulus values with a small jitter for clarity. The reward values of stimuli that share each of the two non-informative features varied from each other due to the design of the reward schedule even though these values were close to 50% on average. (**E**) The error in estimated probabilities (approximation error) based on different learning strategies. Learning about the informative feature and informative conjunction provided the lowest error, whereas learning about the non-informative features and conjunctions lead to a similar error as learning about the informative feature alone (see leftmost point on the blue curve and rightmost point on the red and yellow curves).

Overall, we found that participants performed better than chance (Fig. 1B), and their performances reached steady state after about 150 trials. By examining how well different reinforcement learning models account for participants’ choice behavior, a previous study verified that participants learned about both the informative feature and the conjunction (Farashahi and Soltani, 2021). However, their study did not investigate how participants arrived at this learning strategy, whether some participants deviated from this strategy in a systematic way, and the role of attention in learning. Here, using a combination of model-free analyses and fitting choice data with more complex reinforcement learning models, we characterized how participants’ choice behavior was affected by different attentional strategies, how these attentional strategies interacted with value learning and decision making, and how these interactions affected the overall performance.

### Learning in multidimensional environments is guided by the informative feature and conjunction

To determine how participants adjusted their choices based on reward and choice history early in the experiment, we applied a mixed-effects logistic regression to the first 150 trials of the experiment, before the performance converges (see Materials and Methods for more details). We found that participants’ choices could be significantly predicted by the reward outcome from the previous trial associated with the informative feature (β = 0.19, *SE* = 0.04, *t*(9970) = 4.27, *p* < 0.001; Fig. 2A) and informative conjunction (β = 0.17, *SE* = 0.08, *t*(9970) = 2.15, *p* = 0.03; Fig. 2A). This means that whenever one of the two choice options (stimuli) in the current trial shared the same informative feature or informative conjunction as the chosen stimulus from the previous trial, participants were more likely to choose it if the previous choice was rewarded and avoid it if the previous choice was not rewarded (i.e., win-stay and lose-switch based on feature). In contrast, there was no evidence that participants adjusted their choices according to feedback based on the non-informative features or non-informative conjunctions (*p* > 0.05). These results provide evidence for differential learning about more informative features or conjunctions. In contrast, we found that learning about the non-informative features (including the ones that are constituents of the informative conjunction) and the non-informative conjunctions was attenuated.

**Figure 2.**
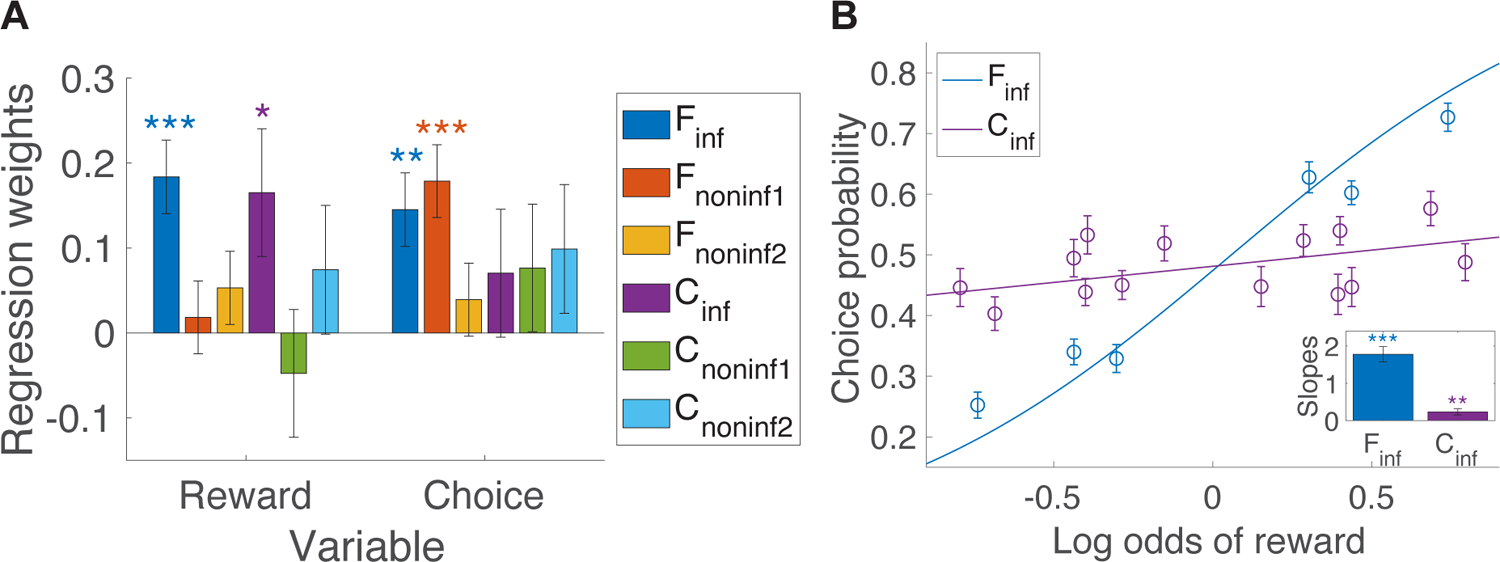
Model-free analysis of reward credit assignment and decision making. (**A**) Analysis of choice behavior during the first 150 trials of the experiment. Plot shows the regression weights from the mixed effects logistic regression model to predict choice using features of choice and reward outcomes in the previous trial. Early choices can be significantly predicted by reward outcome associated with the informative feature and informative conjunction, but not non-informative features or conjunctions.

In addition, we found that participants’ choices were significantly influenced by the previous trial’s choice on the informative feature (β = 0.15, *SE* = 0.05, *t*(9970) = 2.78, *p* = 0.005) and non-informative feature 1 (β = 0.18, *SE* = 0.05, *t*(9970) = 3.57, *p* < 0.001). Overall, participants were more likely to repeat choosing an option that shared either the informative feature, or the non-informative feature 1, with the previously chosen option regardless of the previous outcome. Such choice auto-correlation in general could be explained by a positivity bias in reward learning that weighs positive outcomes more heavily than negative outcomes (Palminteri and Lebreton, 2022). The elevated level of choice auto-correlation in the informative feature and the first non-informative dimensions could also be a consequence of biases in learning or choice, as the effect of positivity bias is amplified through attentional effects on learning. Although the logistic regression analysis cannot separate the influence of attention on choice vs. learning (Katahira, 2018) it enabled us to detect biases in choice behavior that imply differential processing of different features or conjunctions of the stimuli. These results motivated us to investigate the mechanisms through which this attentional bias could have emerged and how it could have interacted with value learning (see the next section).

We also verified that participants’ steady-state choice behavior reflected knowledge about the values of both the informative feature and the informative conjunction. To that end, we used the log ratio of the objective reward probabilities/values of the two options along the informative feature and conjunction dimensions to predict participants’ choice in the last 150 trials using a mixed effects logistic regression model. We found that the log ratio of values along both the informative feature (β = 1.78, *SE* = 0.20, *t*(10047) = 8.71, *p* < 0.001) and the informative conjunction (β = 0.23, *SE* = 0.08, *t*(10047) = 2.89, *p* = 0.003) significantly predicted participants’ steady-state choices, suggesting that participants learned the values of both the informative feature and conjunction and made choices based on a weighted combination of these values as reflected in the best fit logistic curves based on the informative feature and conjunction values (Fig. 2B). We also fit mixed-effects logistic regression models using the objective reward values of both the non-informative conjunctions and the stimulus (the objective values of the non-informative features are all too close to 0.5 to be meaningful). We found that the model that best explained choice behavior considered both the informative feature and the informative conjunction (*X*^2^(4) = 81.73, *p* < 0.001), compared to the model that used only the informative feature, and this model achieved the overall best Akaike Information Criterion (AIC) compared to other models that took into account the non-informative conjunctions’ values, or the stimulus/object values.

Based on the above analyses, we conclude that in the multidimensional reward learning task, participants’ initial behavioral adjustments indicated a higher sensitivity to the reward and choice history of certain features and conjunctions over other ones. This adaptive strategy allowed participants to learn an approximate value representation by learning the values of the informative feature and informative conjunction without having to learn the object/stimulus values directly. This had lasting effects on the participants’ behavior, as they made their choices by combining the values from the informative feature and conjunction once their performances had reached steady state.

Choices can also be significantly predicted by the informative feature and one of the non-informative features of previous choice, but not other variables. One, two, and three asterisks indicate *p* < 0.05, *p* < 0.01, and *p* < 0.001, respectively. (**B**) Analysis of choice during the last 150 trials of the experiment.

Plot shows the fit of late choice data using the informative feature (blue) and conjunction (purple) as indicated in the legend. Inset shows the regression weights from the mixed effects logistic regression analysis of choice, using ground truth reward values of the informative feature and conjunction. Choices later in a session were strongly informed by the values of the informative feature and informative conjunction.

### Attention is guided jointly by the informative feature and conjunction, and only affects learning

The above model-free analyses verified that participants’ credit assignment and/or decision making were biased towards certain features and conjunctions. Next, to explain how these attentional biases emerge and exert their influences, we constructed various reinforcement learning (RL) models that included different attentional mechanisms, and fit choice behavior with these models. The general architecture of the models (Fig. 3A) was inspired by the hierarchical decision making and learning model proposed by Farashahi and colleagues (Farashahi et al., 2017b). In these models, a set of nine feature-encoding units (three for each feature), 27 conjunction-encoding units (nine for each conjunction), and 27 object-encoding units are tuned to different dimensions of the stimuli. Each of these units projects to a corresponding value-encoding unit via synapses that undergo reward-dependent plasticity. This allowed the value-encoding units to estimate reward values associated with individual features, conjunctions of features, and objects. In addition, feature- and conjunction-value encoding units send input to the corresponding attentional-selection circuits that in turn, provide feedback to modulate the gain of stimulus-encoding units. This feedback could modulate decision making by prioritizing values associated with some stimulus dimensions over others. The ensuing attention-weighted value signals drive the decision-making circuit that generates choice on each trial and could result in harvesting reward. The reward outcome on each trial in turn modulates the update of synapses between sensory- and value-encoding units (Soltani and Wang, 2010). In addition to gain modulation during decision making, attention could also differentially modulate the rate of synaptic updates by changing the gain of the pre-synaptic stimuli encoding units, which ultimately modifies the learning rates or associability of different stimuli dimensions (Kruschke, 2001).

**Figure 3.**
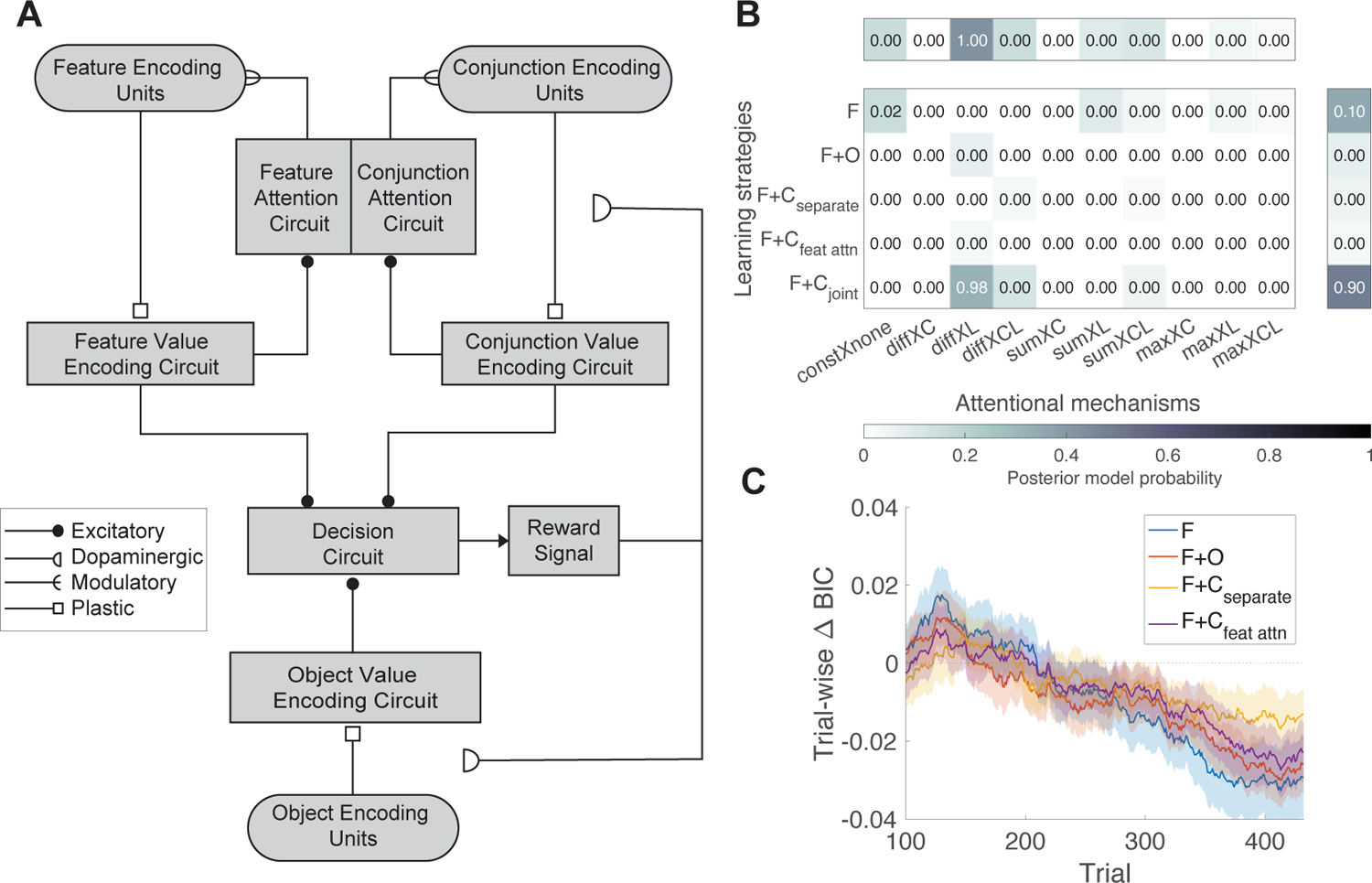
Computational model for learning in high-dimensional environments with value-driven attention. (**A**) Illustration of the model’s architecture. Sensory units encode different features, conjunctions, and object/stimulus identities of the choice options. These units project to feature-value, conjunction-value, and object-value encoding units via plastic synapses that undergo reward-dependent plasticity, allowing the latter units to estimate the corresponding reward values. Feature-value and conjunction-value encoding circuits send feedback to feature and conjunction attention circuits, which potentially interact with each other. Using these value signals, the attention circuit calculates modulatory signals that feedback into the sensory encoding units to modulate the gain of feature/conjunction encoding, which in turn modulates decision making and/or learning. (**B**) Results of random effects Bayesian Model Selection (BMS). Reported values in the middle panel are protected exceedance probability (pxp) of all models. The column and row above and to the right are the results of family-wise BMS aggregating across different types of value learning strategies and across different attentional mechanisms, respectively. The name of attentional mechanisms is given by how attention is calculated (const: constant and uniform, diff: absolute difference, sum: sum / average, max: maximum) and when attention is applied (none: no attentional modulation, C: during choice, L: during learning, CL: during choice and learning). For example, diff X L denotes that attention is calculated based on absolute difference and modulates value updates. (**C**) The average differences in trial-wise BIC between the best model (*F* + *C_joint_*, diff X L) and all other models with the same attentional strategy, but distinct value learning strategies. Shaded areas indicate the SEM. A moving average of window size 100 was applied for visualization purposes, not in hypothesis testing.

To test how learned reward values could drive attention, we compared three possible relationships between these values and attention (in addition to no relationship): summation, absolute difference, and maximum (see **Materials and Methods**). All of these functions could be implemented by canonical recurrent neural network circuits and had been used in prior studies to calculate attention based on subjective reward values, sometimes in much simpler reward schedules where these possibilities could be hard to distinguish from each other (Anderson et al., 2011; Hunt et al., 2014; Niv et al., 2015; Soltani et al., 2016; Farashahi et al., 2017b; Leong et al., 2017; Gluth et al., 2018; Daniel et al., 2020; Farashahi and Soltani, 2021; Pettine et al., 2021; Soltani et al., 2021). In our model, reward values of the two presented options along different dimensions are first passed through one of the above functions, and then normalized across dimensions to have a sum of 1, resulting in one attention weight per feature or conjunction dimension (e.g., color or color/shape conjunction). Attention could modulate decision making, learning, both, or neither. We assumed that during decision making, attention could modulate the relative weights of a stimulus’ feature and conjunction values in determining the log odds of choosing that stimulus. Furthermore, the learning rates of feature and conjunction value updates were modulated by the attention weights to model the effects of attention during learning. We also considered five *learning strategies*: (1) feature-based learning (*F*); (2) feature- and object-based learning (*F* + *O*); (3) feature- and conjunction-based learning with separate attention, where attention weights on the features and conjunction are calculated separately (*F* + *C_separate_*; (4) feature- and conjunction-based learning where attention only biases features, whereas attention is uniform over conjunctions (*F* + *C_feature attn_*); and (5) feature- and conjunction-based learning attention is calculated based on the integrated values of features and conjunctions, modulating both similarly (*F* + *C_joint_*). The last three variations of value learning strategies were tested to investigate possible mechanisms of previously unexplored interaction between attention based on features values and conjunctions values. This comprises 50 models in total.

After fitting the parameters of all models on the trial-by-trial choice behavior of participants using maximum likelihood estimation, we applied Bayesian model selection (BMS) to the Bayesian information criterion (BIC) of all models. The model that best explained our choice data (*F* + *C_joint_ diff X L*, posterior probability = 0.32, *pxp* = 0.98, Fig. 3B) learned both feature and conjunction values. In this model, attention only modulated learning and not decision making. Moreover, attention to conjunctions was linked to attention to features such that the value of a feature was first integrated with the value of the conjunction of the two other features, and then used to guide attention. The resulting attention weights modulated the learning rates of both that feature and conjunction, but not how they were combined during decision making. This means that more attention was allocated to the feature and conjunction pair whose integrated value had the largest absolute difference between the two options. We validated this result by confirming that the best fit model could be identified when compared with models that use the same learning strategy or the same attentional mechanism (**Fig. S1A** and **Fig. S1B**).

Using family-wise BMS by pooling together models that share the same attentional mechanism or the same learning strategies, we confirmed that across all value learning strategies, the best-fitting attentional mechanism modulated only the value updates and not decision making, and this attentional mechanism was controlled by the absolute value difference between the two alternative options (on each trial) along different dimensions (posterior probability = 0.42, *pxp* > 0.99). This reflects a form of biased credit assignment where each outcome was attributed to certain dimension(s) depending on how well value information along that dimension predicted the outcome. This is consistent with previous findings that learning is faster for informative (Farashahi et al., 2017b) and reliable (Mackintosh, 1975) features but extends these findings to learning for informative conjunctions.

Overall, we also found that across all attentional mechanisms, the learning strategies that best fit our data were feature and conjunction learning with separate attention (posterior probability = 0.49, *pxp* = 0.90), closely followed by feature-based learning (posterior probability = 0.35, *pxp* = 0.10). Models with feature-based learning had a good overall fit compared to many of the *F* + *C_joint_* models because the improvement of fit (with additional parameters) in the latter models was not enough to counteract the penalty for having additional parameters captured by BIC. Nonetheless, the winning model’s attentional mechanism (*diff X L*) was uniquely good at explaining choice behavior among all the *F* + *C_joint_* models. By examining the trial-wise BIC (Equation 11) of the models with the best attentional strategy (*diff X L*), we found the difference between the trial-wise BIC of the *F* + *C_joint_* model and the feature-based learning model with the same attentional mechanism decreased over time (β = −0.05, *SE* = 0.01, *t*(28942) = −4.30, *p* < 0.001), and so did the difference between the trial-wise BIC of the *F* + *C*_joint_ with the mixed feature- and object-based learning model (β = −0.04, *SE* = 0.01, *t*(28942) = −3.35, *p* < 0.001). Consistent with previous studies (Farashahi et al., 2017b; Farashahi and Soltani, 2021), this result suggests a transition from feature-based to more conjunction-based learning, without fully transitioning to object-based learning (Fig. 3C), even after considering feature- and conjunction-based attentional modulations. More importantly, attention might play an important role in guiding a gradual transition in learning strategy: after learning about feature values, instead of learning about the values of all conjunctions, attention can be used to select the most informative conjunction for further learning.

Using the maximum likelihood parameter estimates of the best model (**Table S1**, see **Fig. S1C** and **Fig. S1D** for parameter recovery results), we could infer the trial-by-trial subjective attentional weights in the same way that subjective values could be inferred from the RL models that fit choice data the best. Using this approach, we investigated the distribution and dynamics of attention across the informative and non-informative stimulus dimensions. Because attentional weights add up to 1, we treated their distribution as a categorical probability distribution over the dimensions and applied an information-theoretic approach to characterize their dynamic properties with the following observations. First, we found that the entropy of the attentional weights, which is inversely proportional to how focused attention is, decreased throughout the session (β = −0.17, *SE* = 0.03, *t*(67.00) = −5.72, *p* < 0.001; Fig. 4A). This means that attention became more focused as participants learned about reward values associated with stimuli and their features over the time course of the experiment, consistent with previous work (Niv et al., 2015). Second, the Jensen-Shannon divergence (JSD) of attentional weights across consecutive trials, which is proportional to the changes in attention across trials, increased over time (β = 0.06, *SE* = 0.01, *t*(67.00) = 4.77, *p* < 0.001). Even though value learning was a gradual process in our experiment, the model could allow for stimulus-specific, rapid switching of attention across trials by assuming a large inverse temperature in the softmax function used to calculate the normalized attentional weights (see Equation 8). Indeed, we found the estimated value for this quantity to be relatively large (*Median* = 257.88, *IQR* = 410.17 − 70.39; Fig. 5A), making the competition for the control of attention to be close to a hard winner-take-all.

**Figure 4.**
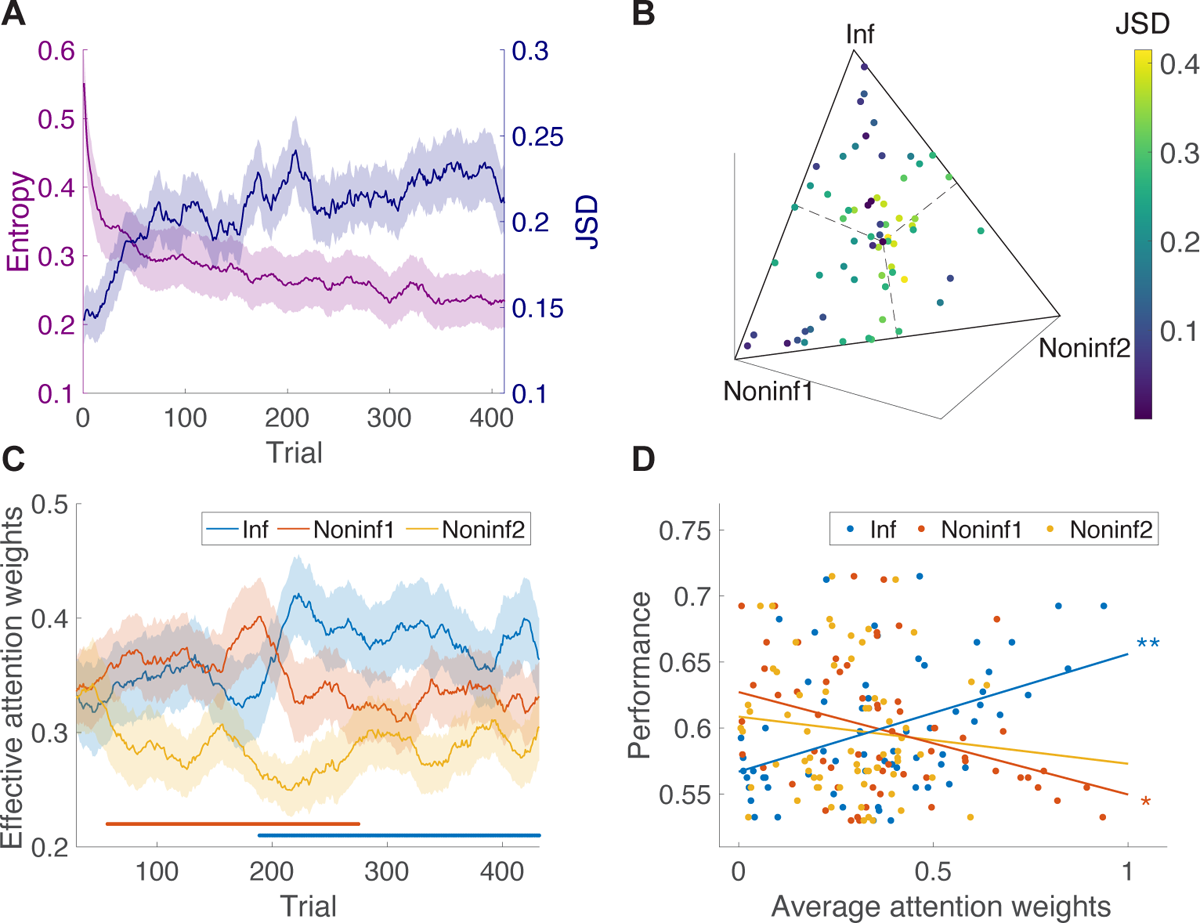
Dynamics of attentional modulation and its effects on performance. (**A**) Plot shows the entropy of the attentional weights and Jensen-Shannon divergence (JSD) between attentional weights in consecutive trials. Entropy rapidly decreased and remained low, suggesting that attention became more focused over time. The increase in JSD suggests that attention tended to switch across trials even late in the session. A moving average with a window size of 20 trials was applied for visualization purposes, not hypothesis testing. (**B**) The average attentional weights for individual participants. The color indicates the average cross-trial JSD for each participant. Participants exhibited a variety of patterns for attentional modulation. Some concentrated on either the informative feature and conjunction pair, or one of the non-informative feature and conjunction pairs (points close to the vertices of the triangle with low JSD). Others oscillated between two or three different dimensions (points far from the vertices with high JSD). (C) Plot shows the time course of smoothed average attentional weights across participants, weighted by participants’ overall sensitivity to reward feedback (β × (α_+_ + α_-_)). The most informative feature and conjunction pair received the most attentional weights on average. A moving average of window size 30 trials was applied for visualization and the cluster-based permutation test. (**D**) Relationship between the allocation of attention and performance. Effective attention weight quantifying credit assignment to the most informative dimensions was associated with increased performance. In contrast, incorrect credit assignment to non-informative dimensions led to poorer performance. One, two, and three asterisks indicate *p* < 0.05, *p* < 0.01, and *p* < 0.001, respectively.

**Figure 5.**
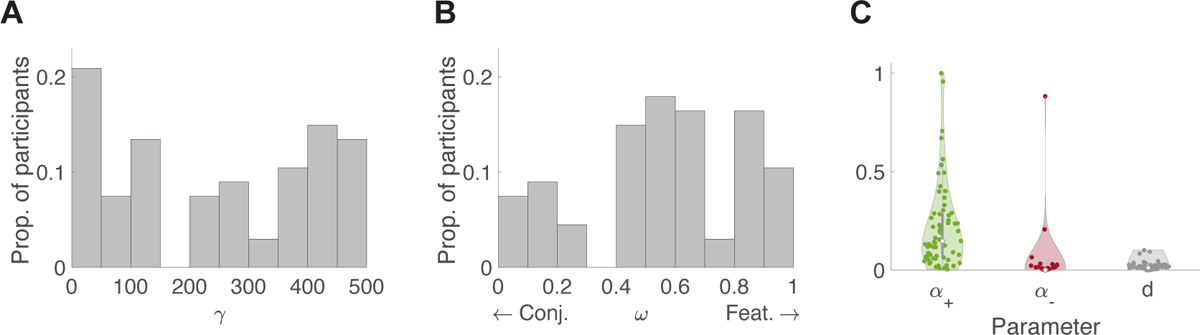
Distribution of key model parameters. (**A**) The distribution of γ values, measuring inverse temperature for attentional selection, in the best-fit model. Higher values of γ correspond to more focused attention. (**B**) The distribution of ω values, measuring relative weighting of feature vs. conjunction for decision making. This distribution suggests that participants tend to consider both feature and conjunction when making decisions but with a slight overall bias towards feature-based learning. (**C**) The distribution of the learning rates for rewarded and unrewarded trials (α_+_ and α_-_) and the decay rate for the value of the unchosen option (*d*). The learning rate for rewarded trials was overall larger than the learning rate for unrewarded trials for most participants, and decay rates were small but non-zero.

Nonetheless, attention was influenced by both feature values and conjunction values and in turn, modulated the learning of both sets of values, as revealed by the distribution of the ω values that quantify the relative influence of feature and conjunction in the model (*Median* = 0.57, *IQR* = 0.81 − 0.45; Fig. 5B).

The distribution of participants’ trial-averaged attentional weights (Fig. 4B) revealed that participants employed a diverse range of attentional strategies. Although some participants focused more on the informative dimensions (i.e., the informative feature and informative conjunction pair), a substantial proportion of them developed attention on the non-informative dimension 1 (the non-informative feature 1 and non-informative conjunction 1). The trial-by-trial, participant-averaged attentional weights also demonstrated an initial bias towards the non-informative dimension 1 (Fig. 4C). To account for each participants’ overall sensitivity to reward feedback, the attention trajectory of each participant was weighted by β × (α_+_ + α_-_) based on their best fit parameters before being averaged, making them a measure of effective attention weights over the entire group. However, the unweighted average of trial-by-trial attention weights (**Fig. S2A**) was used for the following hypothesis testing procedure. Using a one-tailed t-test with a cluster-based permutation test to correct for multiple comparisons (cluster threshold α = 0.05), we discovered a cluster of time points during early trials where more attention was allocated to learning about the non-informative dimension 1 compared to the non-informative dimension 2. As learning progressed, however, more attention was allocated to the informative dimension. A cluster of time points was observed in later trials where more attention was allocated to learning about the informative dimension compared to the non-informative dimension 2. However, we did not find any significant differences between attentional weights to the informative dimension and non-informative dimension 1. It is worth noting that similar attentional weights for the informative feature and non-informative feature 1 does not correspond to similar learning or decision weights for these features, because only the informative feature carries information about reward. One possible explanation for this bias towards non-informative dimension 1 was the asymmetric learning rates (Fig. 5C), which led participants to update their values to a lesser extent after a lack of reward, causing them to learn biased value representations (Cazé and van der Meer, 2013; Katahira, 2018; Palminteri and Lebreton, 2022). This would also explain why the choice history of non-informative feature 1 had a strong effect on participants’ ongoing choices (Fig. 2B). This asymmetry in learning rates was also not simply due to adding the attention component, as it was still present in models without attention (**Fig. S3**). However, its effect could potentially be amplified by attention, leading to differential sensitivity to the choice history associated with different features, as we observed in the current study (Fig. 2A).

Importantly, individual participants’ trial-averaged attention weights predicted their empirical performance such that the more they allocated attention towards learning the values of the informative feature and conjunction, the higher their performance was (ρ(65) = 0.35, *p* = 0.003; Fig. 4D). On the other hand, more attention towards learning the values of non-informative dimension 1 corresponded with lower performance (ρ(65) = −0.27, *p* = 0.03). The correlation between attention towards non-informative dimension 2 and performance was negative but not significant (ρ(65) = −0.13, *p* = 0.30), however, this null result could be due to the small number of subjects who ended up focusing on non-informative dimension 2 (see 4B). Overall, these results suggest that initially, participants tended to develop a bias towards one of the non-informative dimensions. After receiving more reward feedback, however, they transitioned to learning about the correct combination of the informative feature and conjunction. Ultimately, the extent of learning (credit assignment) about the informative dimensions influenced their final performance. These observations highlight the critical role of attention in performing multidimensional learning tasks.

### RL model captures key characteristics of the experimental data

To ensure that the aforementioned results were not due to the best model capturing specific sequence of choices made by the participants and that our best model is able to replicate the main pattern of behavioral results, we simulated the MDPL task using the estimated parameters from the best model (Wilson and Collins, 2019). To that end, we simulated 50 sessions of the experiment using individual participants’ estimated parameters and the exact order of choice options they were presented with. We found that on average, the best model’s performance matched the empirical learning curve (Fig. 6A). Moreover, the model was able to capture individual differences in performance, as there was a high degree of correlation between the empirical performance and the average simulated performance for individual participants (ρ(65) = 0.60, *p* < 0.001) (Fig. 6B).

**Figure 6.**
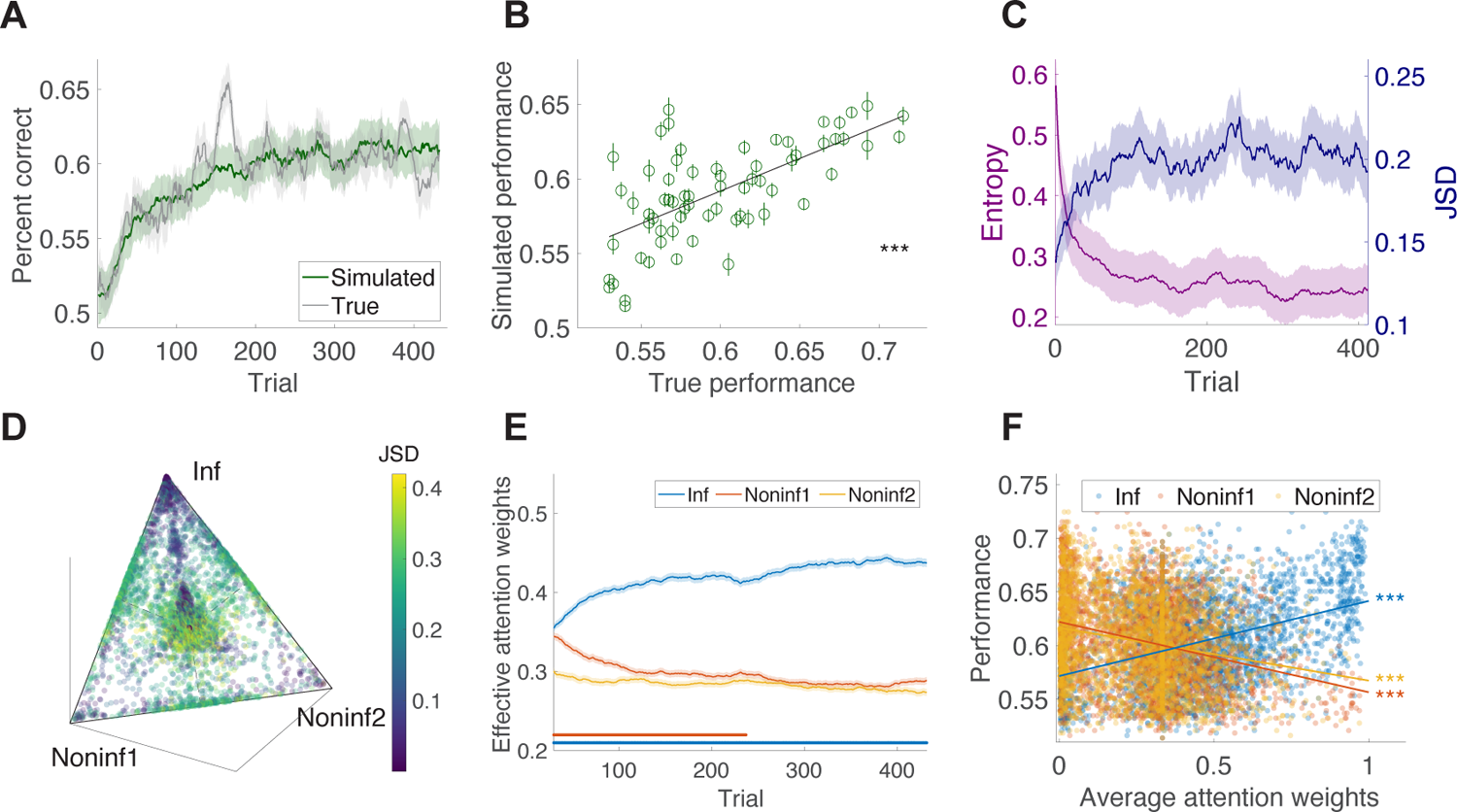
Analyses of simulated data generated by the best model and its estimated parameters for individual participants. (**A–B**) Performance based on simulated data and comparison with observed performance. The simulated performance matched empirical performance on average and across participants. Shades and error bars show the SEM. (**C**) The entropy of the attentional weights and Jensen-Shannon divergence (JSD) between attentional weights on consecutive trials. Simulated trial-by-trial attention became more concentrated over time, and attention can jump sharply across consecutive trials. (D) The average attentional weights for simulated data. Conventions are similar to Fig. 4B. The distribution of attentional weights in simulated data exhibited a diverse set of credit assignment strategies. (E) Simulated trial-by-trial attention. (**F**) Relationship between the allocation of attention and performance in simulated data. Higher attention on the informative feature and conjunction predicted better performance. Triple asterisks indicate *p* < 0.001. Overall, model simulations replicated key aspects of empirical data.

Computing the trial-by-trial simulated attentional weights, we found that attention in the model became more focused over time, as indicated by the decrease of attention weights’ entropy by trial (linear mixed effect model, main effect of trial β = −0.16, *SE* = 0.002, *t*(67.00) = −79.80, *p* < 0.001; Fig. 6C), similar to our experimental observation (Fig. 4A). Moreover, the Jensen-Shannon divergence of attentional weights across consecutive trials increased throughout the trials, suggesting larger jumps of attention across consecutive trials as the session progressed (linear mixed effect model, main effect of trial, β = 0.04, *SE* = 0.001, *t*(67.00) = 36.60, *p* < 0.001; Fig. 6C). In addition, similar to our experimental observations, we found diverse attentional strategies (compare Fig. 6D and Fig. 4B), suggesting that participants’ attentional weights could diverge due to noise in the choice sequence.

We also repeated the cluster-based permutation-test (one-sided t-test, cluster threshold α = 0.05) on simulated data generated by the best model and found that on average, more attention tended to be allocated to the informative dimension than to the second non-informative dimension. In addition, more attention was allocated to non-informative dimension 1 than non-informative dimension 2, but the difference only occurred at the beginning of the session (Fig. 6E, see **Fig. S2B** for the unweighted average attention weights for the simulation). We note that due to the large number of simulated trials for each parameter set, the time course of simulated attentional modulation appears smoother than that of the empirical attention weights (compare Fig. 6E and Fig. 4C). However, the overall amount of stochasticity and cross-trial switches in attention were similar between simulated and empirical data (compare Fig. 6C and Fig. 4A). There was also significant variability in the trajectory of both value functions and attention weights across different runs of the same participant due to stochasticity of choice (**Fig. S4A** and **Fig. S4B**).

Finally, similar to experimental data, we found a significant correlation between performance and attentional strategy in the simulated data such that more attention to the informative dimension was associated with better performance (ρ(3348) = 0.40, *p* < 0.001; Fig. 6F) and more attention towards the non-informative dimensions was associated with lower performance (ρ(3348) = −0.30, *p* < 0.001 for non-informative dimension 1, ρ(3348) = −0.22, *p* < 0.001 for non-informative dimension 2; Fig. 6E).

### Value estimations are biased by the informative feature and conjunction

To test the participant’s representation of the reward contingencies or values more directly, we further analyzed their estimations for reward probabilities associated with the 27 stimuli/objects. Our hypothesis was that the similarities between each participant’s value estimations of different stimuli could be influenced by their attentional biases. For example, a participant who focused their attention on the color dimension would rate all stimuli that share the same color as having similar values. Alternatively, a participant who focused their attention on the conjunction of shape and pattern dimensions would rate all stimuli that share the same shape and pattern configuration similarly but would not rate stimuli that share only a shape or a pattern similarly.

Expanding on methods used in previous work (Farashahi and Soltani, 2021), we fit linear mixed effect models of the participants’ estimation of the reward probability of each stimulus using different reward values based on features, conjunctions, or their combinations as the independent variables. More specifically, we fit five models using the following independent variables: (1) values of the informative feature, *F_inf_*; (2) values of the informative feature and the informative conjunction, *F_inf_*; + *C_inf_*; (3) values of the non-informative feature 1, *C_noninf 1_*; (4) values of the non-informative feature 2, *C_noninf 2_*; and (5) values of stimuli/objects, *O*. Consistent with previous work, we found that the best model for fitting participants’ estimates on estimation trials was a model that predicted value estimates with the values of the informative feature and the informative conjunction as reflected in the adjusted *R*_R_values (Fig. 7B). This model showed the lowest Akaike Information Criterion (AIC) compared to all other models (see the likelihood ratio test comparing to the second-best model that includes only the informative feature in Table 1: *X*^2^(4) > 21.25, *p* < 0.001). Importantly, the worst model was the one that used the stimulus/object values equal to the ground-truth reward probabilities.

**Figure 7.**
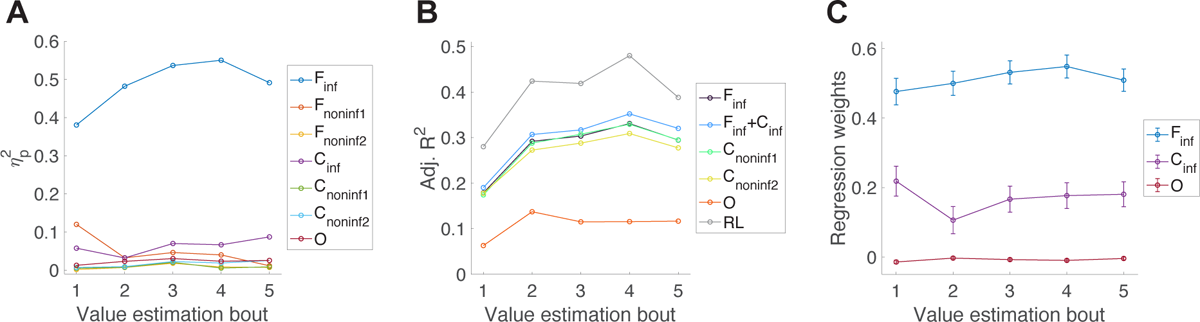
Analyses of reward probability estimates reveal that participants’ estimates were mostly influenced by the informative feature and informative conjunction. (**A**) Partial eta-squared effect sizes (η^2^) from ANOVA. Overall, the reward probability estimates were significantly influenced by the informative feature and the informative conjunction. (**B**) Adjusted *R*^)^ for predicting reward probability estimates in each bout of estimation trials using different objective reward values and estimated subjective reward values using the best RL model. Across all five estimation bouts, subjective reward values based on the best RL provided the best fit followed by a model that included both the objective informative feature and the informative conjunction. (**C**) Regression weight on the marginal probabilities along the informative feature, informative conjunction, and on the ground truth probabilities. Error bars show the SEM.

**Table 1.** Comparison of different reward values’ ability to predict participants’ reward probability estimates across five bouts of estimation trials (*N* = 67). Reported values are the goodness-of-fit metrics (smaller values correspond to better models) based on linear mixed models with different reward values as predictors. The bold values show the best model in each bout, which consistently was the model in which subjective values based on the RL models were used. The underlined values indicate the best model based on objective values calculated based on the reward schedule (as opposed to subjective values estimated by RL models).

|  | Est. bout 1 | Est. bout 2 | Est. bout 3 | Est. bout 4 | Est. bout 5 |
| --- | --- | --- | --- | --- | --- |
| $F_{inf}$ | -LL=2562.07<br>AIC=5136.14<br><u>BIC=5169.14</u> | -LL=2389.29<br>AIC=4790.58<br><u>BIC=4823.59</u> | -LL=2299.19<br>AIC=4610.38<br>BIC=4643.38 | -LL=2262.57<br>AIC=4537.13<br><u>BIC=4570.14</u> | -LL=2231.88<br>AIC=4475.76<br>BIC=4508.76 |
| $F_{inf} + C_{inf}$ | -LL=2551.01<br><u>AIC=5122.01</u><br>BIC=5177.02 | -LL=2378.67<br><u>AIC=4777.33</u><br>BIC=4832.34 | -LL=2288.18<br><u>AIC=4596.35</u><br>BIC=4651.36 | -LL=2248.43<br><u>AIC=4516.87</u><br>BIC=4571.88 | -LL=2212.18<br><u>AIC=4444.36</u><br><u>BIC=4499.36</u> |
| $C_{noninf1}$ | -LL=2565.59<br>AIC=5143.17<br>BIC=5176.18 | -LL=2392.94<br>AIC=4797.89<br>BIC=4830.89 | -LL=2295.46<br>AIC=4602.91<br><u>BIC=4635.91</u> | -LL=2263.92<br>AIC=4539.84<br>BIC=4572.85 | -LL=2231.15<br>AIC=4474.30<br>BIC=4507.30 |
| $C_{noninf2}$ | -LL=2562.44<br>AIC=5136.87<br>BIC=5169.87 | -LL=2406.55<br>AIC=4825.10<br>BIC=4858.10 | -LL=2313.55<br>AIC=4639.10<br>BIC=4672.10 | -LL=2284.04<br>AIC=4580.07<br>BIC=4613.08 | -LL=2246.90<br>AIC=4505.80<br>BIC=4538.81 |
| $O$ | -LL=2651.50<br>AIC=5315.00<br>BIC=5348.00 | -LL=2523.29<br>AIC=5058.57<br>BIC=5091.58 | -LL=2463.17<br>AIC=4938.35<br>BIC=4971.35 | -LL=2453.61<br>AIC=4919.22<br>BIC=4952.22 | -LL=2386.02<br>AIC=4784.03<br>BIC=4817.03 |
| $RL$ | -LL=2457.04<br>**AIC=4926.08**<br>**BIC=4959.08** | -LL=2233.19<br>**AIC=4478.38**<br>**BIC=4511.38** | -LL=2158.97<br>**AIC=4329.95**<br>**BIC=4362.95** | -LL=2071.04<br>**AIC=4154.07**<br>**BIC=4187.08** | -LL=2127.15<br>**AIC=4266.30**<br>**BIC=4299.30** |

In order to test whether participants learned about stimulus/object values in addition to feature and conjunction values, we fit mixed-effect models using reward values of the informative feature and the informative conjunction, as well as the reward values of the stimuli/objects. We found the regression weight on the informative feature to be always significantly bigger than 0 (β > 0.48, *p* < 0.001 for all five bouts; Fig. 7C; **Table 2**), and this was also the case for the weight on the informative conjunction (β > 0.11, *p* < 0.007 for all five bouts; Fig. 7C). The regression weight on the stimulus/object values was either non-significant or significant but negative, suggesting that participants did not meaningfully learn about the reward values of objects other than those explained by the informative feature and conjunction (**Table 2**). These are consistent with our results based on the analysis of choice behavior, suggesting that participants employed an efficient learning strategy using the informative feature and conjunction.

**Table 2.** Coefficients from the linear mixed-effects modeling of the participants’ reward probability estimates (*N* = 67). The independent variables were the log odds of reward values for the informative feature, informative conjunction, and the stimuli/object, with coefficients denoted as *F_inf_*, *C_inf_*, and *O*, respectively.

| Bout | Name | Coefficient | SE | p-value |
| --- | --- | --- | --- | --- |
| 1 | $F_{inf}$ | 0.48 | 0.04 | <0.001 |
| | $C_{inf}$ | 0.22 | 0.04 | <0.001 |
| | $O$ | -0.01 | 0.005 | 0.004 |
| 2 | $F_{inf}$ | 0.5 | 0.03 | <0.001 |
| | $C_{inf}$ | 0.11 | 0.04 | 0.007 |
| | $O$ | 0 | 0.004 | 0.504 |
| 3 | $F_{inf}$ | 0.53 | 0.03 | <0.001 |
| | $C_{inf}$ | 0.17 | 0.04 | <0.001 |
| | $O$ | -0.01 | 0.004 | 0.09 |
| 4 | $F_{inf}$ | 0.55 | 0.03 | <0.001 |
| | $C_{inf}$ | 0.18 | 0.04 | <0.001 |
|  | *O* | -0.01 | 0.004 | 0.024 |
| 5 | *F<sub>inf</sub>* | 0.51 | 0.03 | <0.001 |
|  | *C<sub>inf</sub>* | 0.18 | 0.04 | <0.001 |
|  | *O* | 0 | 0.004 | 0.324 |

Although the above analyses revealed that participants acquired knowledge about the ground-truth reward values in terms of the informative feature and conjunction, they cannot detect if they exhibited deviations from the ground truth that could reflect attentional biases. Such biases can be uncovered by fitting ANOVA models that use the features of each stimulus and their interactions to predict the participants’ reward probability estimates, and comparing the variance explained by each feature and their interactions. This method does not depend on the ground truth reward values (calculated from the reward schedule) and therefore, is more capable of capturing biases in learning. Using this method we found that, consistent with the previous method, both the informative feature and the interaction of the two non-informative features (the informative conjunction) explained a significant amount of variance in the value estimates from all estimation trials (informative feature: *F*(2,528) > 31.40, *p* < 0.001, η^2^ > 0.38; informative conjunction: *F*(4,528) > 2.71, *p* ≤ 0.03, η^2^ > 0.03). We performed the same analysis on each of five estimation bouts separately and found that non-informative feature 1 explained a significant amount of variance in the first bout of value estimates despite the fact that this feature carried no information (*F*(2,528) = 7.15, *p* = 0.001, η^2^ = 0.12; **Table 3**). Consistent with the observation in the previous sections (Fig. 2B and Fig. 4C), this suggests a sub-optimal attentional focus on the non-informative feature 1 dimension at the beginning of the session. We speculate that this happened due to the small amount of information carried by this dimension, the noise in the choice process, the bias induced by asymmetric learning rates, and fluctuations in attention during learning. Nonetheless, later in the session, the informative feature and the informative conjunction explained most variance in value estimates (**Table 3**), consistent with the results from RL modeling that attention became more focused on the informative feature and conjunction after more trials (Fig. 4C).

**Table 3.** Result from the mixed-effects ANOVA analysis of participants’ reward probability estimates (*N* = 67).

|  | Est. bout 1 | Est. bout 2 | Est. bout 3 | Est. bout 4 | Est. bout 5 |
| --- | --- | --- | --- | --- | --- |
| $F_{inf}$ | $F(2, 528)=31.40$<br>$p<0.001$<br>$\eta_p^2=0.38$ | $F(2, 528)=47.01$<br>$p<0.001$<br>$\eta_p^2=0.48$ | $F(2, 528)=43.69$<br>$p<0.001$<br>$\eta_p^2=0.54$ | $F(2, 528)=50.22$<br>$p<0.001$<br>$\eta_p^2=0.55$ | $F(2, 528)=58.25$<br>$p<0.001$<br>$\eta_p^2=0.49$ |
| $C_{noninf1}$ | $F(2, 528)=7.15$<br>$p=0.001$<br>$\eta_p^2=0.12$ | $F(2, 528)=2.08$<br>$p=0.13$<br>$\eta_p^2=0.03$ | $F(2, 528)=3.06$<br>$p=0.05$<br>$\eta_p^2=0.05$ | $F(2, 528)=2.29$<br>$p=0.11$<br>$\eta_p^2=0.04$ | $F(2, 528)=0.97$<br>$p=0.38$<br>$\eta_p^2=0.01$ |
| $C_{noninf1}$ | $F(2, 528)=0.13$<br>$p=0.88$<br>$\eta_p^2=0.00$ | $F(2, 528)=0.66$<br>$p=0.52$<br>$\eta_p^2=0.01$ | $F(2, 528)=1.57$<br>$p=0.21$<br>$\eta_p^2=0.02$ | $F(2, 528)=0.72$<br>$p=0.79$<br>$\eta_p^2=0.01$ | $F(2, 528)=0.81$<br>$p=0.44$<br>$\eta_p^2=0.01$ |
| $C_{inf}$ | $F(4, 528)=5.00$<br>$p<0.001$<br>$\eta_p^2=0.06$ | $F(4, 528)=2.71$<br>$p=0.03$<br>$\eta_p^2=0.03$ | $F(4, 528)=6.39$<br>$p<0.001$<br>$\eta_p^2=0.07$ | $F(4, 528)=5.53$<br>$p<0.001$<br>$\eta_p^2=0.07$ | $F(4, 528)=9.09$<br>$p<0.001$<br>$\eta_p^2=0.09$ |
| $C_{noninf1}$ | $F(4, 528)=0.49$<br>$p=0.74$<br>$\eta_p^2=0.00$ | $F(4, 528)=0.79$<br>$p=0.53$<br>$\eta_p^2=0.01$ | $F(4, 528)=1.93$<br>$p=0.11$<br>$\eta_p^2=0.02$ | $F(4, 528)=0.64$<br>$p=0.63$<br>$\eta_p^2=0.01$ | $F(4, 528)=0.91$<br>$p=0.46$<br>$\eta_p^2=0.01$ |
| $C_{noninf1}$ | $F(4, 528)=0.71$<br>$p=0.59$<br>$\eta_p^2=0.01$ | $F(4, 528)=0.89$<br>$p=0.47$<br>$\eta_p^2=0.01$ | $F(4, 528)=2.67$<br>$p=0.03$<br>$\eta_p^2=0.02$ | $F(4, 528)=1.89$<br>$p=0.11$<br>$\eta_p^2=0.02$ | $F(4, 528)=3.27$<br>$p=0.01$<br>$\eta_p^2=0.03$ |
| $O$ | $F(8, 528)=0.85$<br>$p=0.56$<br>$\eta_p^2=0.01$ | $F(8, 528)=1.56$<br>$p=0.13$<br>$\eta_p^2=0.02$ | $F(8, 528)=2.06$<br>$p=0.04$<br>$\eta_p^2=0.03$ | $F(8, 528)=1.58$<br>$p=0.13$<br>$\eta_p^2=0.02$ | $F(8, 528)=1.72$<br>$p=0.09$<br>$\eta_p^2=0.03$ |

To further relate reward probability estimates from the estimation bouts to the RL models used to fit choice data, we also utilized subjective values from the RL models to predict reward probability estimates. To that end, we first computed a weighted average of the subjective values along different feature and conjunction dimensions to compute reward value of stimuli before each estimation bout (i.e., after choice trials 86, 173, 259, 346, 432). As mentioned above, the RL model that best explained participants’ choice data estimated the reward values of features and conjunction and had uniform attention during choice (see Fig. 3B and **Eq. 5**). We z-scored these subjective values from RL modeling for each participant because the inverse temperature parameter allowed these values to assume very different ranges. Using a linear mixed-effects model, we then fit a scaling parameter and an intercept in order to predict the estimated reward probabilities reported by participants using the subjective values from the RL model. We found that reward probability estimates were better fit by subjective values based on the best RL model than by the objective reward values (**Table 1**). This result further illustrates that the winning RL model captured the learning and integration of values along different dimensions well.

Overall, analyses of the participants’ value estimates confirmed our analyses of choice data suggesting that participants learned about the informative feature and the informative conjunction and prioritized learning about these dimensions over other dimensions. In addition, these analyses showed that sub-optimal attentional strategy also led to biases in value estimation. Finally, the fact that subjective probability estimates were well captured by the subjective values based on the best RL model, even though that RL model was not fit on the value estimates in estimation trials, further validates the suitability of this model in capturing behavioral data.

## Discussion

Learning in naturalistic environments relies on the interaction between multiple forms of reward learning, decision making, and selective attention. To investigate this interaction and underlying neural mechanisms, we analyzed behavioral data from a multidimensional probabilistic reward learning task with multiple approaches. Using model-free methods, we found differences in sensitivity to both the reward and choice history associated with different features and conjunctions. Specifically, participants adjusted their behavior by preferentially associating reward outcomes to the informative feature and conjunction of selected stimuli. They also repeated choosing the options that shared the same informative feature (and in some participants, one of the non-informative features) as in the preceding trial. Similarly, their value estimations resembled a weighted combination of the reward values of the informative feature and the informative conjunctions, with an initial bias towards one of the non-informative features that diminished over time.

To explain these observations, we constructed multiple RL models that learned feature, conjunction, and/or stimulus values with different value-based attentional mechanisms that modulated decision making and/or learning. The best model in terms of explaining participants’ choices was the one that learned both feature and conjunction values and deployed attention mainly during learning. In this model, attention was controlled by the difference in reward values of the two options in terms of feature and conjunction and in turn, modulated learning but not choice behavior. Interestingly, feature and conjunction values influenced attention in a cooperative manner such that the values of a feature were first integrated with the values of the conjunction of the two other features, and the resulting attention weights modulated the learning of both feature and conjunction values. RL models allowed us to examine the participant-specific trial-by-trial learning strategies. Specifically, the trial-wise fit of choice data revealed a transition from feature-based learning to mixed feature- and conjunction-based learning. In addition, the amount each participant attended to the informative feature and conjunction correlated with their performance in the task.

Flexibilities in stimuli representation for value learning, either by deploying selective attention to reduce dimensionality (Niv et al., 2015; Leong et al., 2017; Mack et al., 2020) or by acquiring conjunctive representations to increase dimensionality (Rigotti et al., 2013; Bernardi et al., 2020), are two aspects of representation learning (Niv, 2019; Radulescu et al., 2019a, 2021), which have been studied using variety of tasks in different fields. This includes the information integration and weather prediction tasks in category learning (Kruschke, 2001; Ashby and Maddox, 2005), multi-cue conditioning tasks in classical conditioning (Mackintosh, 1975; Pearce and Hall, 1980; Dayan et al., 2000; O’Reilly and Rudy, 2001; Harris, 2006), and variants of multidimensional reinforcement learning (RL) tasks (Niv et al., 2015; Farashahi et al., 2017b; Leong et al., 2017; Farashahi and Soltani, 2021). All these tasks require learning of stimulus-outcome contingencies where the ground truth contingencies contain certain structures to allow the use and generalization of learning strategies. For example, some features provide little or no information about reward outcomes and can be filtered out through selective attention to enable more efficient learning with little impact on performance (Niv et al., 2015; Leong et al., 2017; Mack et al., 2020). Moreover, associations between certain stimulus features and outcomes could be generalizable or context specific (Kruschke, 2001; Ashby and Maddox, 2005), making outcomes predictable based on elemental or configural stimulus representation, respectively (Mackintosh, 1975; Harris, 2006), and these translate to feature-based or object-based learning in RL (Farashahi et al., 2017b). Here, by carefully parameterizing the relationship between the combinations of stimulus features and reward outcome probabilities associated with each three-dimensional choice options, we were able to control the generalizability of the reward schedule to include novel structures that have not been tested in previous studies: i.e., the presence of both an informative feature and an informative conjunction.

Most previous studies on interactions between reinforcement learning and selective attention did not consider the possibility of reward-predictive conjunctions despite evidence that in multidimensional environments, healthy individuals are capable of learning the values of conjunctions/configurations of features (O’Reilly and Rudy, 2001; Farashahi et al., 2017b; Duncan et al., 2018; Ballard et al., 2019; Pelletier and Fellows, 2019). This could be because those studies on configural learning were performed in the context of two-feature choice options with little ambiguity about the informative conjunctions. In contrast, our results based on three-dimensional choice options suggest that in naturalistic environments in which reward values cannot be generalized across features, selective attention can be more complex than a simple competition among features (or conjunctions of features). Instead, value representations based on both features and conjunctions interact to determine attention, which in turn shapes efficient state representations upon which learning can happen. Importantly, this value-guided attention provides an additional mechanism for controlling the tradeoff between adaptability and precision (Farashahi et al., 2017a, 2017b). By focusing on the most task-relevant aspects of the environment, this strategy allows for the conservation of cognitive resources (via selective attention) while developing more sophisticated state-space representations (with conjunctive representations).

The attentional modulation present in the best-fitting model is similar to the process in a hierarchical decision making and learning model that used the relative informativeness of stimulus values and integrated feature values to arbitrate between feature-based and object-based learning, but without a need for attention (Farashahi et al., 2017b). In low-dimensional environments where there are only a few features and stimuli to learn about, feature-based attention may not be required. In contrast, in high-dimensional environments in which there are multiple ways for representing reward values of stimuli (feature-based, conjunction-based, object-based representations, or a mixture thereof), additional adaptive mechanisms are needed. This includes selection between different ways for representing stimulus values such that the resulting representations are parsimonious while that the associated values provide a strong signal to differentiate between choice options.

Attention to reward-predictive features can be instantiated by mixtures of expert models using biologically plausible learning mechanisms (Badre and Frank, 2012; Frank and Badre, 2012; Collins and Frank, 2013; Lee et al., 2014; Cortese et al., 2021). In these models, multiple “expert” modules try to predict the reward outcome, and (approximate) Bayesian inference is used to arbitrate among these modules. In the context of multidimensional reinforcement learning tasks, each expert module effectively instantiates a hypothesis about the latent structure of reward contingencies. The observed interaction between feature-based and conjunction-based learning can be implemented using a mixture of experts. However, this requires many independent circuits to perform exact mixture-of-expert inferences. In contrast, our proposed models assume that the value learning circuit itself can exhibit attentional biases due to internal activity within the decision making circuit (Szabo et al., 2006; Pannunzi et al., 2012) without having a dedicated hypothesis testing module.

Interestingly, stochastic, reward-dependent Hebbian synaptic plasticity provides a biologically plausible mechanism for implementing the Naive Bayes algorithm (Soltani and Wang, 2010; Murphy, 2012). Naïve Bayes is an efficient classification algorithm for problems where the different features are conditionally independent given the outcome (Ng and Jordan, 2001; Murphy, 2012). However, when this assumption is violated, as in our experiment in which the conjunction of two features provided additional information about reward probabilities, Naïve Bayes in its simplest form (similar to elemental or feature-based learning) leads to suboptimal behavior. The attentional selection of the informative conjunction during learning is, however, conceptually similar to the hierarchical Naive Bayes that has been proposed to deal with the problem of conditional dependence between features (Han et al., 2005; Langseth and Nielsen, 2006). Similar to our best-fitting model, instead of trying to filter out irrelevant features or irrelevant conjunctions, hierarchical Naive Bayes selects features to create informative conjunctions. Although the models tested here were not motivated by optimality principles, the connection between our findings and above algorithms may provide a normative explanation for why this attentional mechanism is adopted.

Although we assumed that value-based attention could affect both choice and learning (but only found evidence for attentional effects on learning), other theoretical works have suggested that attention at choice and learning could serve different roles to improve decision making in high dimensional and uncertain environments (Dayan et al., 2000), and that the outcome of the choice may play a role in switching attention before value updates (Kruschke, 2001). There is also evidence for different degrees of attentional modulation at choice and learning, and for outcome-dependent attention during learning (Kruschke, 2001; Akaishi et al., 2016). In the models tested here, the switching of attention was dependent on gradual value learning but not on instantaneous reward feedback. Due to concerns of parameter identifiability, we also did not test for different attentional mechanisms during choice and learning. Eye-tracking could be used as a model-independent measure of attention to avoid this issue (Leong et al., 2017). Although uniform attention to multiple features individually might be hard to differentiate from attention to the conjunctions of these features even with eye-tracking, this method could provide auxiliary data for fitting attention modulated RL models (Radulescu et al., 2019b).

Due to difficulty of learning the values of 27 stimuli, we focused on learning with a stable reward schedule. However, naturalistic environments can be volatile, with changing reward contingencies. These changes usually do not manifest as randomly reassigning reward probabilities across all stimuli/objects, but as localized changes within a task-relevant dimension or changes in the identity of the task relevant dimension. These two types of changes mirror the intra- and extra-dimensional shifts in the set-shifting literature (Owen et al., 1991) and may have distinct effects on learning and choice behavior. The presence of reversals has consequences on learning and choice behavior in both humans and non-human primates (Behrens et al., 2007; Farashahi et al., 2017a, 2019; Soltani and Izquierdo, 2019). Similarly, the contributions of attention to learning may be easier to detect when there are reversals. This is because attention at learning would lead to blocking of learning when the informative dimension is changed and to facilitation of learning when there is a reversal along the previously informative dimension.

Attention at choice alone would not have these effects on learning but may enhance the effect of blocking and facilitation when acting in conjunction with attention at learning. Although previous studies on multidimensional reward learning have utilized both types of reversals, their effects were not systematically investigated in the context of probabilistic reward learning, and it would be an interesting direction for future research (Kruschke, 2001; Niv et al., 2015; Akaishi et al., 2016; Farashahi et al., 2017b).

Overall, our study provides evidence for the existence and possible origin of attentional modulations in multidimensional reward learning. Nonetheless, specific neural mechanisms by which attention modulates learning are not currently known. For example, although a study found neural evidence for selective reward credit assignment to task-relevant variables (Donahue and Lee, 2015), the relevant variable did not change across sessions in that study (e.g., color was always the reward-predictive cue) and no dynamic interaction between learning and attention was necessary to solve the task. Future studies could investigate how attentional effects emerge with value learning, and how it in turn modulates learning, possibly through feedback connections involving multiple brain regions (Roelfsema et al., 2010).

## Acknowledgment

We would like to thank Chanc Orzell for helpful comments on the manuscript. This work was supported by National Science Foundation CAREER Award (BCS1943767) to A.S.

## Supplementary Information

**Supplementary Table S1.** Summary statistics of estimated parameters of the best-fitting model.

| Name | 25% Percentile | Median | 75% Percentile |
| --- | --- | --- | --- |
| *bias* | -0.07 | 0.05 | 0.21 |
| $\beta$ | 19.11 | 24.60 | 42.00 |
| $\omega$ | 0.45 | 0.57 | 0.81 |
| $d$ | 0.01 | 0.01 | 0.02 |
| $\alpha_+$ | 0.07 | 0.15 | 0.29 |
| $\alpha_-$ | 0.00 | 0.00 | 0.01 |
| $\gamma$ | 70.39 | 257.88 | 410.17 |

**Supplementary Figure S1.**
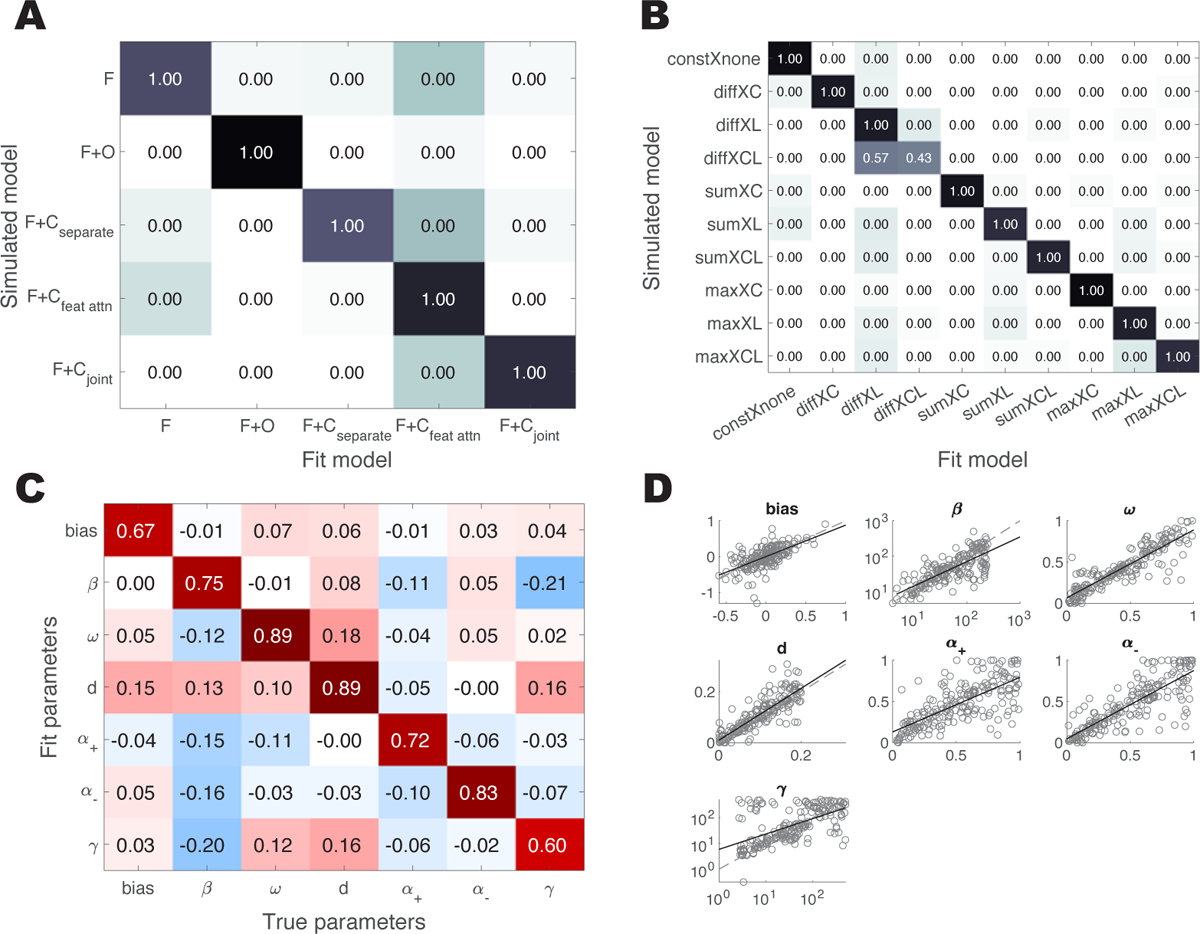
Model and parameter recovery. (A–B) Using Bayesian Model Selection, we compared the winning model with other models with: the same attentional mechanism (modulation of learning based on value difference) and different learning strategies (A), and the same learning strategy (F+C learning with joint attention) but different attentional mechanisms (B). The reported values are pxp, and the color of the cells show the posterior model probabilities. The true models were all well-recovered with pxp=1, except the models with value-difference-based attention modulating both choice and learning were hard to recover from the model where only learning was modulated. However, the false model did not fit the data significantly better than the true model (Wilcoxon’s signed-rank test, Δ*BIC* = 1.20, *p* = 0.83). (C–D) Parameter recovery of the winning model calculated as the Spearman’s correlation between true and estimated parameters is shown in D. All estimated parameters were significantly correlated with the true parameters (Spearman’s ρ > 0.60, *p* < 0.001), and the correlations between different parameters were small.

**Supplementary Figure S2.**
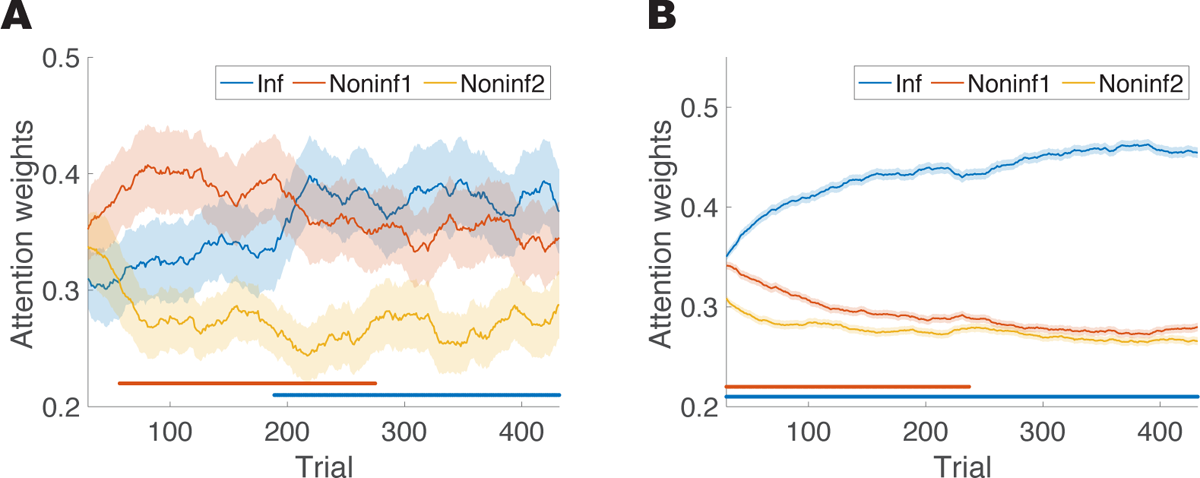
Trial-by-trial average attentional weights, not weighted by the product of inverse temperature and learning rate, during (**A**) empirical and (**B**) simulated choice sequences. The difference between the informative and first non-informative dimensions was diminished, but the same qualitative pattern was still present.

**Supplementary Figure S3.**
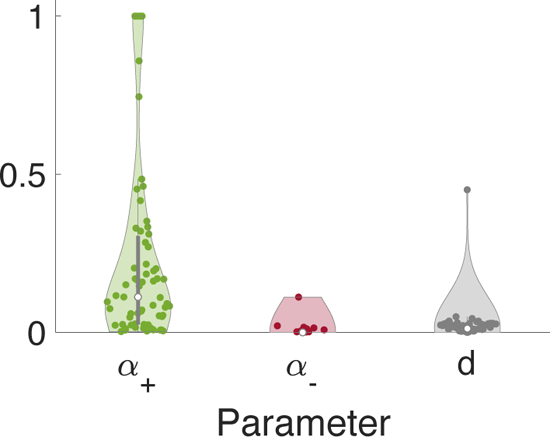
Estimated learning rates for the feature-based and conjunction-based learning in the model without attention. Learning rates were larger for rewarded than unrewarded trials (positivity bias). This suggests that the positivity bias in learning rate was not due to the inclusion of attention or the inadequacy of the current attentional mechanisms in explaining learning from a lack of reward.

**Supplementary Figure S4.**
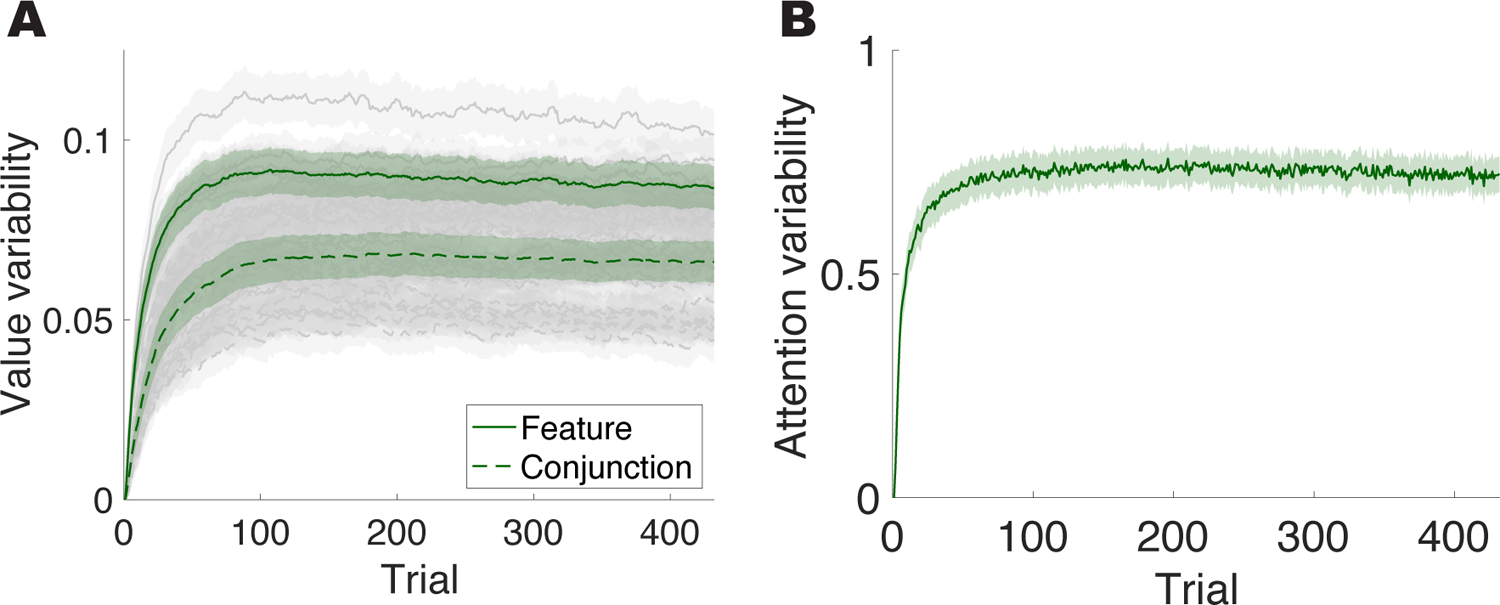
Variability of value estimations and attention weights across simulations of each individual participant for the winning model. (A) The variability of each feature/conjunction value, as quantified by the standard deviation of each value across different simulation runs of the same set of empirical parameters. The grey lines are for each individual value and the dark green lines are aggregated across all feature and conjunction values. This shows that even with the same set of parameters, due to stochasticity in choice, variability in subjective values can persist throughout the session. (B) The variability in attention, as quantified by the KL-divergence between attention weights in each simulation run and the average attention weights across all simulation runs, for each subject. Similar to the subjective values, variability in attention persists throughout the session.

